# Modern Japanese ancestry-derived variants revealed the formation process of the current Japanese regional gradations

**DOI:** 10.1101/2020.12.07.414037

**Authors:** Yusuke Watanabe, Jun Ohashi

**Affiliations:** Department of Biological Sciences, Graduate School of Science, The University of Tokyo, Tokyo 113-0033, Japan; Genome Medical Science Project Toyama Project, National Center for Global Health and Medicine, Tokyo 162-8655, Japan

**Keywords:** admixture, ancestry-derived variants, Japanese population history, Jomon, phenotypic variation

## Abstract

Modern Japanese have two major ancestral populations: the indigenous Jomon hunter gatherers and continental East Asian farmers. To figure out the formation process of current Japanese population, we developed a reference-free detection method of variants derived from ancestral populations using a summary statistic, the ancestry-marker index (*AMI*). We confirmed by computer simulations that *AMI* can detect ancestry-derived variants even in an admixed population of recently diverged source populations with high accuracy, which cannot be achieved by the most widely used statistics, S*, for identifying archaic ancestry. We applied the *AMI* to modern Japanese samples and identified 208,648 single nucleotide polymorphisms (SNPs) that were likely derived from the Jomon people (Jomon-derived variants). The analysis of Jomon-derived variants in 10,842 modern Japanese individuals recruited from all over Japan revealed that the admixture proportions of the Jomon people varied between prefectures, probably due to the differences of population sizes of immigrants in the final Jomon to the Yayoi period. The estimated allele frequencies of genome-wide SNPs in the ancestral populations of modern Japanese suggested their phenotypic characteristics possibly for adaptation to their respective livelihoods; higher triglycerides and blood sugar for the Jomon ancestry and higher C-reactive protein and eosinophil counts for continental ancestry. According to our findings, we propose a formation model of modern Japanese population; regional variations in admixture proportions of the Jomon people and continental East Asians formed genotypic and phenotypic gradations of current Japanese archipelago populations.

## Introduction

Modern Japanese populations are composed with three main populations: the Ainu, who live mainly in Hokkaido; the Ryukyuan, who live mainly in Okinawa; and mainland Japanese, who live in Honshu, Shikoku, and Kyushu (Figure S1). A now-established theory of the formation processes of Japanese populations, a dual structure model, was proposed based on morphological findings by Hanihara 1991^1^. This model assumes that Japanese originated through a mixture of the Jomon people, the Neolithic hunter-gatherers who settled in the Japanese archipelago during the Jomon period (from 16,500 years before present (YBP) to 2,800 YBP)^2–4^, and the immigrants came to the Japanese archipelago with their rice-farming technology from continental East Asia around the beginning of the Yayoi period (around 2,800 YBP)^4^. The rice farming of continental immigrants subsequently spread throughout Japan and brought about a major transformation in ancient Japanese society. According to the dual structure model, compared to mainland Japanese, the Ainu and the Ryukyuan were genetically less influenced by immigrants. Genetic studies not only supported the dual structure model, but also revealed the detailed population history of Japanese archipelago^5–11^. Whole-genome analyses extracted from the remains of the Jomon people showed that the Jomon people were highly differentiated from other East Asians, forming a basal lineage to the East and Northeast Asians^8,10,11^. The genetic relationship between a Jomon individual and other East Asians suggested that the ancestral population of the Jomon people is one of the earliest-wave migrants who might have taken a coastal route on the way from Southeast Asia toward East Asia^11^. It was also revealed that the Jomon people were genetically closely related to the Ainu/Ryukyuan, and that 10–20% of genomic components found in mainland Japanese are derived from the Jomon people^8,10^. Recent studies found that, in addition to the “East Asian” population which is closely related to modern Han Chinese, the “Northeast Asian” population also contributed to the ancestry of modern Japanese^12,13^. Cooke et al 2021 showed the deep divergence of the Jomon people from continental populations including the “East Asians” and “Northeast Asians”, so it can be concluded that the modern mainland Japanese are a population with genomic components derived from a basal East Asian lineage (i.e., the Jomon people) and from continental East Asians. We collectively refer the two continental ancestral populations pointed out in Cooke et al. as “the continental East Asians” in this article, unless a distinction is necessary.

The dual structure model is a powerful hypothesis about the formation history of the Japanese archipelago populations. However, there is still a mystery that is not fully explained by the long-established dual structure model; the formation process of regional variations in the Japanese populations. Several previous studies pointed out the east-west variation of morphological traits and classical genetic markers, and Hanihara referred to these studies that “The differences today between east and west Japan likely originated in the Jomon and Yayoi ages.”^1^ More recently, some studies have demonstrated the genomic regional variation among Japanese archipelago populations at the genome-wide scale^5,6,14,15^. In particular, based on large collections of Japanese samples from across the Japanese archipelago, previous studies showed that the Tohoku, Kanto, and Kyushu populations are genetically more closely related to the Ryukyuan, while the Kinki and Shikoku populations are more closely related to continental East Asians^14,15^. There also seems to be the regional gradations in the genetic factors that define the phenotype of the Japanese. Isshiki et al. found that the polygenic score (PS) for height was higher in the mainland Japanese who were more closely related to the Han Chinese^16^. Thus, regional differences in genetic and phenotypic characteristics exist among the Japanese, which are speculated to be defined by the “degree of genetic relatedness to the Han Chinese”. We can summarize these conjectures by Hanihara and later anthropologists into the following hypotheses; the genetic regional differences among the modern mainland Japanese are caused by regional geographical differences in the admixture proportion of the Jomon and immigrants from continental East Asia, dating in the Late Jomon to Yayoi periods. To verify this hypothesis, we focused on the “Jomon-derived variants” in the modern Japanese archipelago population.

In populations derived from a mixture of two source populations, recombination between haplotypes from different source populations inevitably occurs after the admixture event. As a result, haplotypes from two ancestral populations are patchily present in the chromosomes of admixed population, and the alleles in the haplotypes from each ancestral population are expected to be in linkage disequilibrium (LD) with each other. In this study, we developed a method using a summary statistic, the Ancestry-Marker Index (*AMI*), to detect ancestry-marker variants derived from the Jomon people (i.e. Jomon-derived variants) in modern mainland Japanese. A key feature of the *AMI* is that it does not require genomes obtained from Jomon skeletal specimens. The *AMI* was developed with inspiration from S*, detecting archaic-hominin-derived haplotypes using the specific single nucleotide polymorphisms (SNPs) in the out-of-Africa population which assumed to be originated from admixture of archaic hominins and early Eurasians^17–20^. Since the Jomon people were highly differentiated from other East Asians^8,10^, they are expected to have had specific variants that are not found in present-day East Asian populations other than the Japanese. Thus, it is likely that modern mainland Japanese also have specific variants derived from the Jomon people. The *AMI* detects the Jomon-derived variants based on the LD between Japanese specific variants. We could successfully extract the Jomon-derived variants from the real genomic data of the Japanese. We conducted comprehensive analyses using Jomon-derived variants as proxies for the magnitude of Jomon ancestry and genetic markers for estimations of polygenic traits of modern Japanese ancestry (a mixture of Jomon and continental ancestry) which was not evident from morphological characteristics of skeletal remains, which enabled us to elucidate the mechanisms by which genetic and phenotypic regional gradations among the Japanese arose. Based on our findings, we propose a model of formation process of current regional populations in the Japanese archipelago.

## Results

### Development of the *AMI*, a summary statistic to detect ancestry-derived variants

First of all, we attempted to detect the Jomon-derived genomic components by S*^17,18^, one of the most widely used statistics for identifying archaic ancestry, and found that S* was unable to detect the Jomon-derived genomic components by a coalescent simulation assuming population history of modern Japanese (Figure S2 - S4 and Supplemental information). The Jomon population has been isolated in the Japanese archipelago for a much shorter period since its divergence from other Asian populations than the divergence time of archaic Hominin and modern humans, which is the main target of S*, so there seem no haplotypes in which Jomon-specific variation is in proximity being enough to S* analysis. We therefore developed a new summary statistic, the *AMI*, to distinguish Jomon-derived variants (type 1) from other types of Japanese specific variants; (type 2) variants derived from continental East Asians and (type 3) novel variants in Japanese lineages after admixture (Figure 1). The *AMI* was developed based on the concept that the Jomon people, a part of the ancestral population of modern Japanese, were highly genetically differentiated from other continental East Asian populations, and modern Japanese have inherited specific variants accumulated in Jomon lineages which are not observed in other East Asian populations (type 1; Jomon-derived variants in Figure 1). Our first step is extracting specific variants of modern Japanese to remove variants that have emerged in the non-Jomon ancestry lineage after the divergence of the Jomon people and the non-Jomon ancestry (i.e., orange triangle in Figure 1). In the next step, we calculate *AMI* to distinguish Jomon-derived variants from other types of Japanese specific variants. There are two types of Japanese specific variants other than (type 1) Jomon-derived variantsz; (type 2) variants derived from continental East Asians, which emerged in the continental East Asian lineages and were moved into the Japanese lineages through the admixture, but were eventually lost in the East Asian population; and (type 3) novel variants that emerged only in Japanese lineages after the admixture (Figure 1). Of these Japanese specific variants, the Jomon-derived variants (type 1) are considered to be accumulated on the same haplotype or to be in LD with each other. In other words, variants belonging to (type 1) are expected to have more variant pairs in LD compared to (type 2 and 3). To calculate the *AMI*, we first compute the LD coefficient *r*^2^ between Japanese specific variant pairs. The *AMI* is the count of variants where *r*^2^ exceeds the cutoff for a focal Japanese specific variant, divided by the density of specific variants per kb. Calculating the *AMI*, those in LD with more Japanese specific variants will be distinguished as Jomon-derived variants (type 1). To confirm the usefulness of the *AMI*, we performed a coalescent simulation assuming a mixture of the Jomon people and continental East Asians (Figure S2). In the 1 Mb simulation, Japanese specific variants (types 1, 2, and 3) were extracted from each genealogy, and calculated the *AMI* for each Japanese specific variant. The distributions of *AMI* showed that Jomon-derived variants (type 1) had larger *AMI* values than the other Japanese specific variants (types 2 and 3) (Figure 2A). The receiver operating characteristic (ROC) analysis showed that Jomon-derived variants (type 1) could be distinguished from the other Japanese specific variants (types 2 and 3) by the *AMI* (area under the curve [AUC] = 0.91; Figure 2B). The Youden index, a measure of the cutoff value, was 28.0374. We performed ROC analyses, varying the *r*^2^ threshold from 0 to 0.8 and calculated AUC values to examine differences in detection ability due to differences in *r*^2^. We found that (type 1) cannot be detected at 0, that there is almost no difference in detection ability between 0.01 and 0.2, and that the detection ability decreases as *r*^2^ exceeds 0.2 (Figure S5). We performed further simulations, varying the split time between the Jomon people and continental East Asians, the effective population size, or Jomon ancestry proportion in the current Japanese population to confirm the robustness of *AMI* to different population histories. Though the value of the Youden index varied depending on the population history assumed, the Jomon-derived variants could be accurately detected (Figure S6).

**Figure 1.**
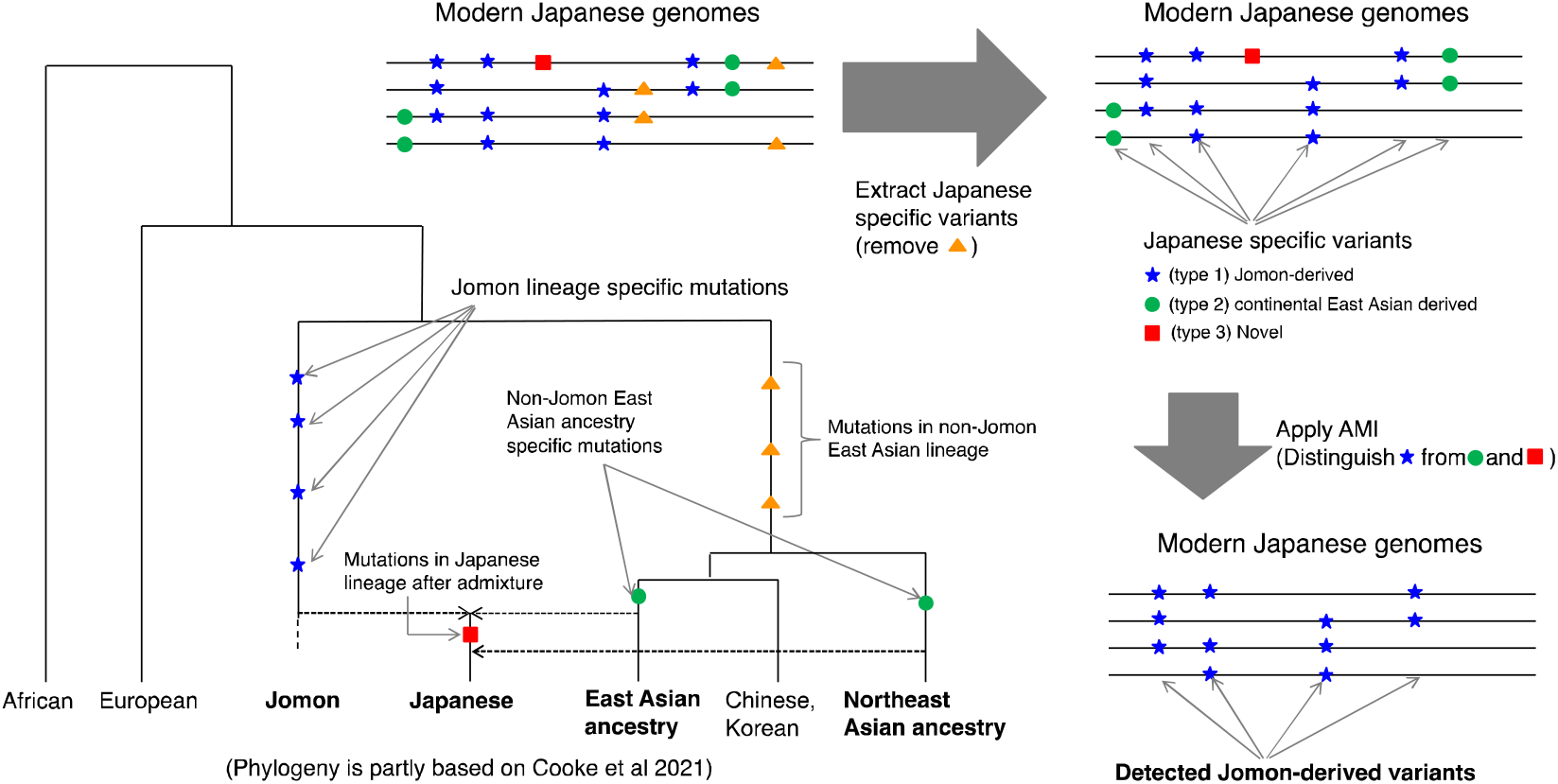
Overview of our *AMI*-based detection method of Jomon-derived variants of modern Japanese. Phylogenetic relationships between modern Japanese, the Jomon people and continental East Asians^4–7^. Modern Japanese genomes contain a mixture of variants that occurred on each ancestral lineage of modern Japanese (Jomon and continental East Asian ancestries including “East Asians” and “Northeast Asians” in Cooke et al., 2021). The Jomon people were highly differentiated from continental East Asians, so modern Japanese genomes have Jomon-lineage specific variants (blue stars in the phylogeny) that are not found in other modern continental East Asians (Chinese, Koreans and so on); *AMI* is an index to distinguish Jomon-derived variants from other Japanese-specific variants based on the linkage disequilibrium status.

**Figure 2.**
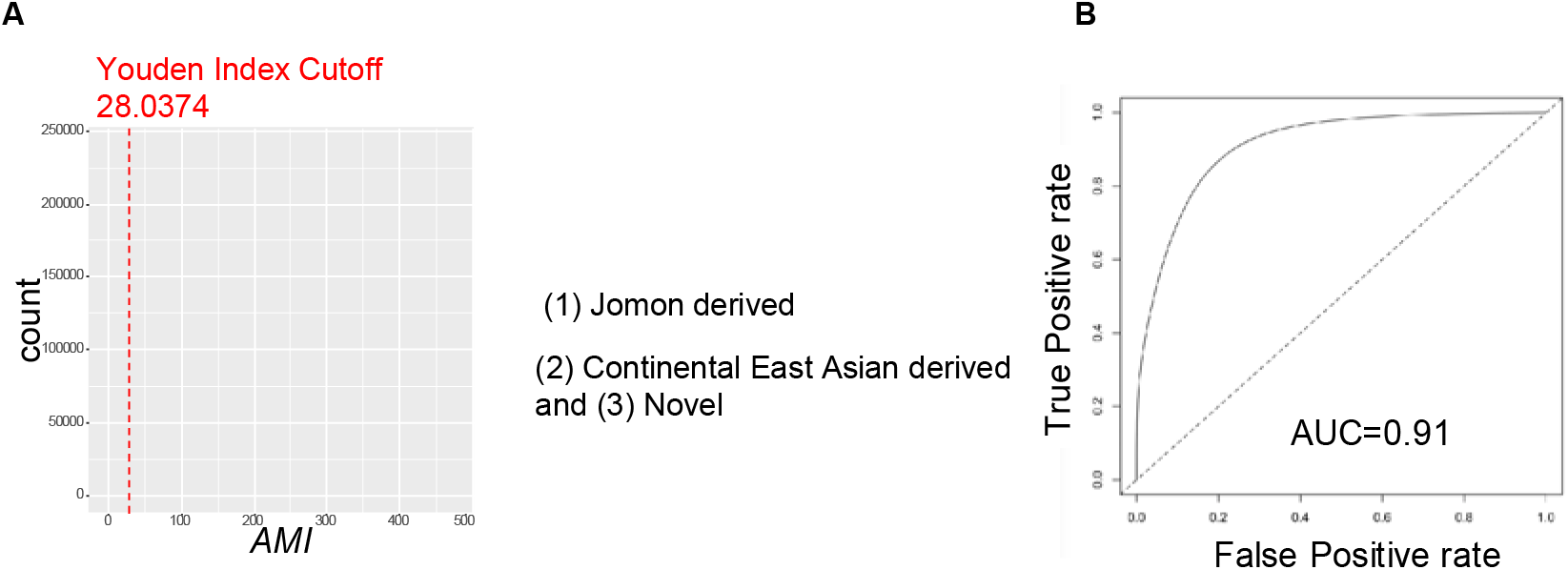
Performance of the *AMI* for the detection of the Jomon-derived variants. (A) Distribution of *AMI* simulated by msprime. The histogram of *AMI* for the Jomon-derived variants (type 1) and the other variants (types 2 and 3) are shown. The red dashed line indicates the threshold of *AMI* (28.0374) obtained from ROC analysis for the detection of the Jomon-derived variants (type 1). (B) ROC curve illustrating the performance of the *AMI* for the detection of the Jomon-derived variants. The ROC curve was drawn based on the simulated data shown in Figure 2A. The *AMI* showed high accuracy (AUC = 0.91) for discriminating the Jomon-derived variants (type 1) from the other variants (types 2 and 3).

### Detection of Jomon variants in real data

We focused only on bi-allelic SNPs to detect Jomon-derived variants. Using the dataset of 87 Korean Personal Genome Project (KPGP) Koreans^21^ and 26 global populations of 1000 Genome Project (1KG)^22^, approximately 1.7 million SNPs were found to be specific to mainland Japanese (1KG JPT). Of these 1.7 million SNPs, 208,648 SNPs exceeding the threshold of *AMI* were regarded as Jomon-derived variants. Jomon-derived variants were distributed throughout the autosomal genome (Figure S7).

To examine the detection accuracy of Jomon-derived variants, we computed the Jomon allele score (*JAS*), calculated based on Jomon-derived vatiant count, for two Jomon individuals, Ikawazu^9,11^ and Funadomari^10^, and 104 mainland Japanese. If Jomon-derived variants were properly detected by the *AMI*, the *JAS* of the Ikawazu or Funadomari Jomon were expected to be higher than those of mainland Japanese. Of the JPT mainland Japanese, NA18976 was genetically close to continental East Asians in principal component analysis (PCA) (Figure S8) and was expected to have a lower *JAS*. The distribution of the *JAS* is shown in Figure S9. The mean *JAS* of 103 mainland Japanese individuals, excluding NA18976, was 0.0164. As expected, NA18976 had the lowest *JAS*, 0.00269, which was much lower than that of the other mainland Japanese. The *JAS* of Ikawazu Jomon and Funadomari Jomon were 0.0523 and 0.0555, respectively, indicating that the Jomon-derived variants were found more frequently in Jomon people than in the modern mainland Japanese. These results suggest that the *AMI* could detect SNPs derived from the Jomon people. It should also be noted that the *JAS* values were only a few percent even for both the Ikawazu and Funadomari Jomon individuals, which suggests that the number of Jomon-specific variants obtained from *AMI* analyses of modern Japanese were several tens of times greater than that obtained from the whole genome sequence of a single Jomon individual.

### Detection of regional genetic differences in mainland Japanese by Jomon-derived variants *JAS* by region and prefecture

To verify our hypothesis about regional variation in mainland Japanese, we calculated the average *JAS* for each geographic region and prefecture from imputed genotypes of 3,917 Jomon-derived variants of 10,842 Japanese individuals previously used for regional population genetic analysis^15^. We removed the Hokkaido samples, which were largely affected by the immigration of Japanese after the Meiji period (1,868~), for subsequent analysis and a total of 10,412 samples were used. The samples of each prefecture except for Hokkaido were divided into ten regions: Tohoku, Kanto, Hokuriku, Chubu, Tokai, Kinki, Chugoku, Shikoku, Kyushu, and Okinawa in accordance with a previous study^23^ (Figure S1 and Table S1). The *JASs* in these ten geographical regions are presented in Figure 3A and Table S2. We found that the *JAS* was the highest in Okinawa (0.0255), followed by Tohoku (0.0189) and Kanto (0.018), and the lowest in Kinki (0.0163), followed by Shikoku (0.016). In prefecture scale, the average *JAS* in mainland Japan tended to be higher in prefectures located in the northernmost and southernmost parts of mainland Japan (Figure 3B and Table S3). The *JAS* was especially high in Aomori (0.0192), Iwate (0.0195), Fukushima (0.0187), and Akita (0.0186) prefectures of the Tohoku region, as well as Kagoshima Prefecture (0.0186) in Kyushu. Interestingly, the *JAS* in Shimane Prefecture (0.0184) was on the same level as the prefectures in Tohoku and Kagoshima in Kyushu, being consistent with the genetic affinity in the Izumo individuals to Okinawa and Kyushu individuals in Jinam et al^24^. Japanese individuals in these prefectures are considered to possess more Jomon-derived genomic components than those in other prefectures. Prefectures with lower *JASs* were in the Kinki and Shikoku regions, including Wakayama (0.0157), Nara (0.0156), Kochi (0.016), Tokushima (0.0161), and Mie (0.0161). These populations are considered to have more genomic components derived from continental East Asians. The *JAS* of each prefecture and the principal component 1 (PC1) value, which was obtained from the PCA of a previous study by the allele frequency of autosomal 183,708 SNPs in each prefecture^15^ are plotted in Figure 3C. The *JAS* was strongly correlated with PC1 (*R* = 0.91, two-sided *t*-test *P* = 2.2 × 10^−16^). The geographic distribution was not changed by tighter cutoff values (*AMI* > 100) for the detection of Jomon-derived variants by *AMI* (Figure S10 and SI).

**Figure 3.**
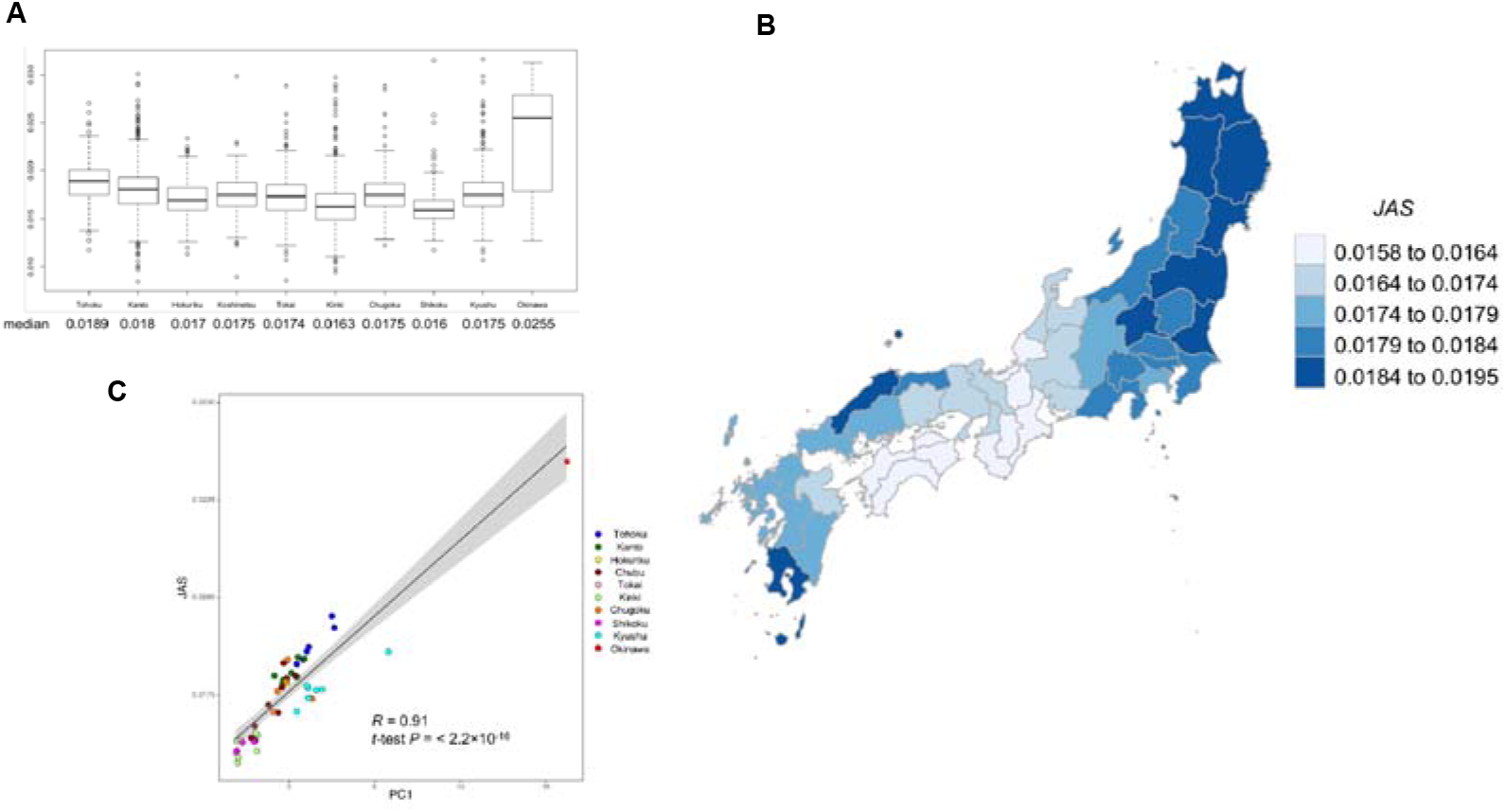
JAS of each region of Japan. (A) Distribution of *JAS* in ten regions. The boxplot of the *JAS* is presented for each of the ten regions, excluding Hokkaido. (B) *JAS* of each prefecture in mainland Japan. The average *JAS* by prefecture was calculated. Hokkaido and Okinawa prefectures are not illustrated. The prefecture with the higher average *JAS* is illustrated by the darker color. (C) Relationship between the PC1 of the PCA performed in a previous study by the allele frequency of autosomal 183,708 SNPs in each prefecture and average *JAS*. Each prefecture was colored according to the region of Japan in Figure S1. Horizontal axe: PC1, vertical axe: average *JAS*. Pearson’s correlation coefficients (*R*), *P* values, regression lines and 95% CI are shown.

We assumed that the regional differences in the *JAS* were related to regional differences in population size during the Jomon period, so we examined the correlation between *JAS* and three indexes related to the Jomon population size. The *JAS* for each region was significantly correlated with several archaeological population indices (Figure S11). The positive correlation between *JAS* and population size in each region (Figures S11A and S11B) and the negative correlation between *JAS* and population growth rate occurring from the Late Jomon period to the Yayoi period (Figure S11C) suggests that the smaller the population size in the Jomon period, the lower *JAS* in modern mainland Japan (i.e., the higher contribution of genomic components of immigrants from continental East Asia). Furthermore, we referred to a previous study that estimated the timing of the arrival of rice farming by Bayesian techniques based on the radiocarbon dating of charred rice remains by constructing two different models a and b^25^. They suggested that after rice farming arrived in northern Kyushu, it reached the Kinki and Shikoku regions earlier than southern Kyushu, which is highly consistent with the low level of *JAS* in the Kinki and Shikoku regions. The relationship between *JAS* and the estimated arrival dates of rice farming in each region suggested that the lower the *JAS*, the earlier the arrival of rice farming tended to be (Figure S12; *R* = −0.71, two-sided *t*-test *P* = 0.05 for model a; *R* = −0.67, two-sided *t*-test *P* = 0.071 for model b). In summary, we can conclude that genetic gradations among the regional population of modern Japanese were mainly caused by differences in the admixture proportion of the Jomon people, perhaps due to population size differences in each region during the Final Jomon to the Yayoi period.

### Allele frequencies estimation by Jomon-derived haplotypes reveals the phenotypic characteristics of the Jomon and continental East Asian ancestry

We devised an estimation method of the allele frequencies of genome-wide SNPs in the Jomon people, the ancestors of modern Japanese, prior to their admixture with continental populations without using Jomon individual genomes. Modern Japanese haplotypes surrounding a focal SNP can be classified into “Jomon-derived haplotypes” and “continental haplotypes” according to the presence Jomon-derived variants (Figure S13). The allele frequency of the Jomon people in the focal SNP can be estimated by the proportion of each allele within Jomon-derived haplotypes (Figure S13). The proportion of each allele within continental haplotypes is the allele frequency of the continental ancestry of modern Japanese. Using 419 modern Japanese genomes (Tokyo Healthy Control, THC dataset)^26^, we applied our method for 6,481,773 SNPs in the Jomon and continental ancestries of modern Japanese (“THC Jomon ancestry” and “THC continental ancestry”) to estimate the allele frequencies. The allele frequencies of THC ancestries were verified by ancient genomes previously reported (11 Jomon and three Kofun individuals^9–11,13^) and genomes of THC modern Japanese and Han Chinese (CHB)^22^ (Figure 4). Kofun individuals, who were excavated from mainland Japan 1,300-1,400 years ago, have similar Jomon ancestry proportion of modern Japanese^13^. First, a pairwise *f3* assuming the Yoruba people as an outgroup was used to test whether the allele frequencies of THC Jomon ancestry resembles that of the actual Jomon individuals (Figure 4). The *f3* value of the THC Jomon ancestry with each Jomon individual from previous studies took higher values (*f3* > 0.04) than that of with Kofun, modern Japanese and modern Chinese. The *f3* values calculated between the THC Jomon ancestry and the Jomon individuals were comparable to that of the Jomon individual pairs (Figure 4). These results indicate that the THC Jomon ancestry were genetically close to the actual Jomon individuals, and that we can successfully infer allele frequencies of the Jomon people by the Jomon-derived haplotypes found in modern Japanese. These results also provide strong assurance that the Jomon-derived variants of modern Japanese we extracted were indeed originated through the Jomon people.

**Figure 4.**
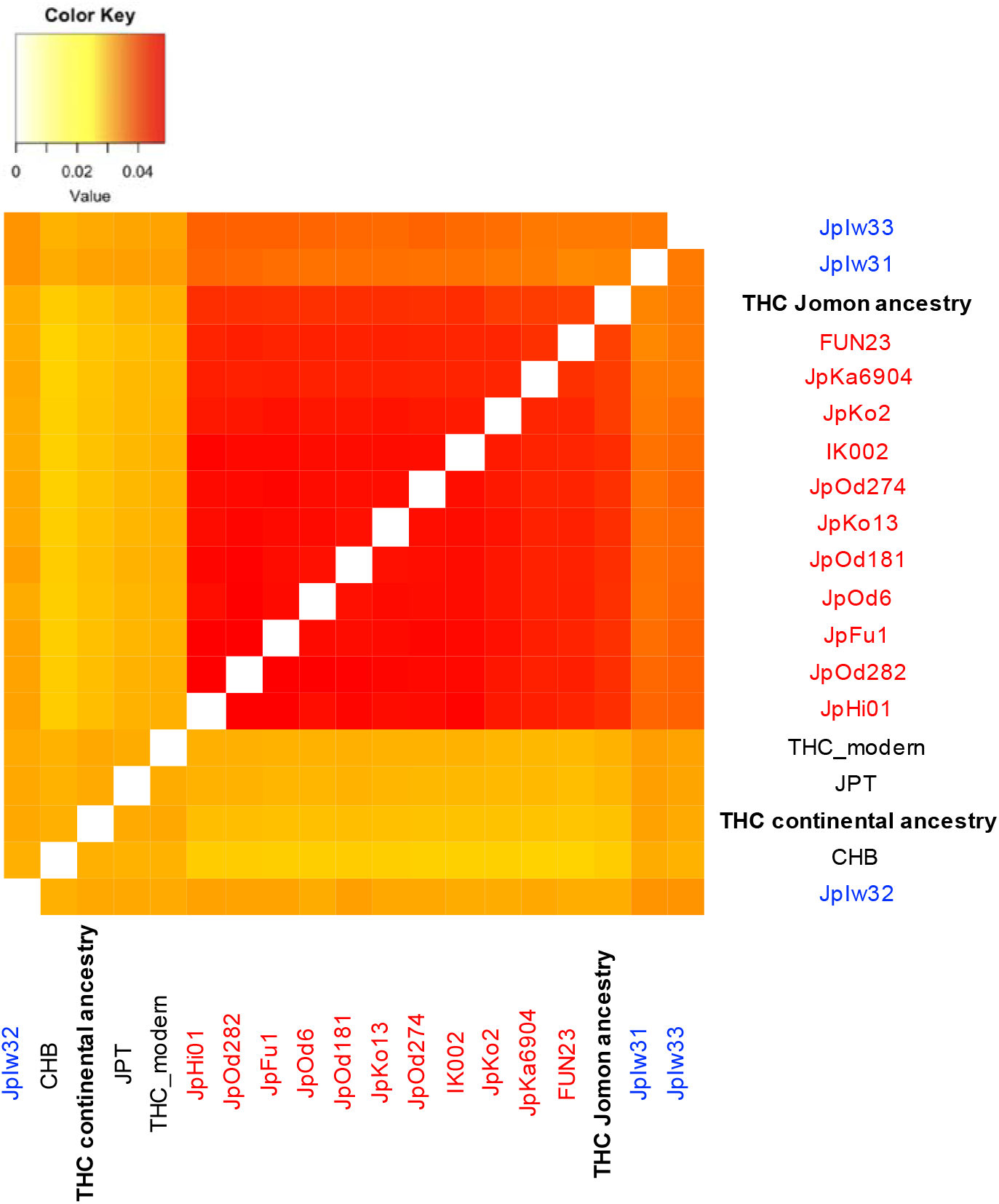
Accuracy of the allele frequency estimation in the Jomon people from Jomon-derived haplotypes of modern Japanese. A heatmap of pairwise f3 statistics between the actual Jomon (red) and Kofun (blue) individuals, modern Japanese (JPT and THC), modern Han Chinese (CHB) and Jomon/continental ancestries of modern Japanese inferred from Jomon-derived haplotypes of modern Japanese.

To estimate the phenotypic characteristics of the Japanese ancestral populations, we combined previous Genome-wide association study (GWAS) results^27–29^ with allele frequencies of genome-wide SNPs of THC modern Japanese, THC Jomon and continental ancestries. We calculated the mean 2*βf* value for each population allele frequencies. The *2βf* is the 2 × (effect allele frequency) multiplied by (effect size) for each SNP with GWAS *P* value lower than the threshold values we set (*P* = 0.01, *P* = 0.001 and *P* = 0.0001), and the mean 2*βf* over genome-wide SNPs allows us to evaluate the average phenotype in a given population for the trait. The mean 2*βf* corresponds to the average polygenic score (PS)^30^ within a focal population. We calculated the mean 2*βf* values for 60 traits in the previous quantitative trait locus (QTL) GWAS (Table S4 and S5). First focusing on THC modern Japanese, the mean 2*βf* values for all 60 traits in the modern Japanese were close to the continental ancestry. Deviation of the population frequency from the ancestry proportion in an admixed population is known to be a signal of positive natural selection after the admixture event^31,32^. The mean 2*βf* values of THC modern Japanese reflected the high percentage of continental ancestry in the Japanese population (80~90%), and it seems unlikely that phenotypes of the Jomon ancestry became dominant in the modern Japanese due to natural selection. Next, to identify traits with particularly large differences between Jomon and continental ancestries, we calculated *D* statistics based on the null distribution obtained from simulations. We can infer that the larger the absolute value of *D*, the more significant the phenotypic difference between Jomon and continental ancestries were. *D* takes a positive value when the mean 2*βf* is greater for THC Jomon ancestries than for THC continental ancestries. The *D* values for each trait, varying the GWAS *P* value threshold, are shown in Figure 5A - 5C and Table S5. The traits that show significant *D* values differ slightly depending on the GWAS *P* value threshold. This may because when the *P* value threshold is set strictly, the polygenic effect of SNPs with relatively small effect size are eliminated, and a few SNPs with higher effect size may be emphasized. For example, for triglycerides (trait ID: TG), which has the highest *D* value when we set the strict *P* value threshold (Figure 5B and 5C), there is rs964184 C/G on the 3’ UTR of the *ZPR1* gene on chromosome 11 that significantly increases TG (β= 0.16, *P* = 1.4 ×10^−272^), which remarkably differs in allele frequency between THC Jomon ancestry and in THC modern Japanese/THC continental ancestry (94%, 28% and 18%, respectively). We then focus on the following traits for which significant *D* values were obtained in Figure 5A - 5C; triglycerides (trait ID: TG), blood sugar (trait ID: BS) for the positive *D* values and height (trait ID: Height), C-reactive protein (trait ID: CRP), eosinophil count (trait ID: Eosino) for the negative *D* values. These *D* values inferred that Jomon ancestry had genetically shorter stature, higher triglyceride, blood sugar, while the continental ancestry had genetically taller stature, higher CRP and eosinophil counts. With regard to height, our results were very convincing because several previous morphological studies suggested that the Jomon people had shorter statue than the people after the migration from continental East Asia, such as the people of the Yayoi and the Kofun period^33–35^. Concerning the other traits that significantly differ between THC Jomon and continental ancestry, those with positive *D* values appear to be related to nutritional status, while those with negative *D* values appear to be related to resistance to infectious diseases. These phenotypic characteristics seemed to have been genetic adaptations to their respective livelihoods; the Jomon people may have needed to maintain high triglyceride and blood sugar levels in their hunter-gatherer lifestyle, while continental East Asian populations may have needed to increase their resistance to infectious diseases during their agricultural lifestyles. We will provide more detailed description in the Discussion section.

**Figure 5.**
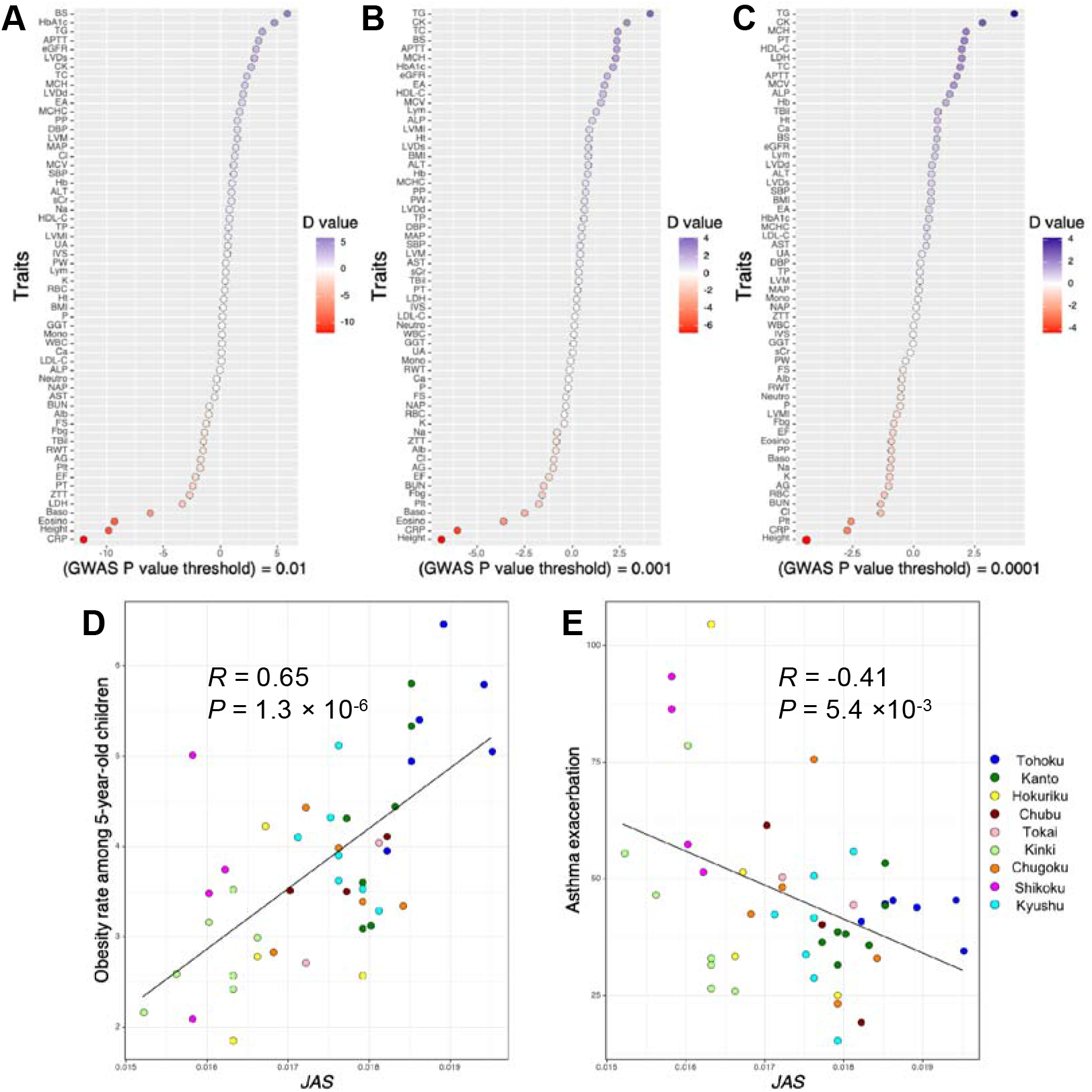
Inference of phenotypic characteristics of the Jomon and continental ancestries and impact of regional admixture proportion differences on regional phenotypic variations of modern Japanese. (A) *D* value for each phenotype, plotted in dark purple when the Jomon ancestry had higher mean 2*βf* values than the continental ancestry, and dark red vice versa. (B) - (D) Histograms of the mean 2*βf* obtained by simulations for height, eosinophil count and blood sugar. Mean 2*βf* was shown for THC modern Japanese (a black line), THC Jomon (a blue line) and continental ancestries (an orange line) for each phenotype. (E) Relationship between the *JAS* and incidence rates of asthma exacerbation of prefectural populations of modern Japanese. Pearson’s correlation coefficients (*R*), *P* values are shown in each figure. Each prefecture is colored according to the region in Figure S1.

Isshiki et al. suggested that PS for height correlated with the mean height of each prefectural population in Japan and those genetically closely related to the Han Chinese have smaller PS for height among regional populations in modern Japan^16^. Combining Isshiki et al. with our finding that the continental ancestry had genetically taller stature, we can strongly argue that the regional variation in height among the Japanese archipelago was caused by differences in the ancestry proportions of the Jomon people, who had genetic factors for shorter statue, and continental East Asians, who had genetic factors for taller statue. In addition to height variations between the regional populations of the Japanese archipelago, we report cases in which regional differences in ancestry proportions affected the regional diversity in a modern Japanese phenotype in relation to triglycerides/blood sugar and eosinophil counts. We focused on regional differences in the obesity rate among 5-year-old children, which is related to triglycerides and blood sugar, and the incidence rates of asthma exacerbation, which is related to eosinophils^36^, among mainland Japanese populations. Eosinophils had strongly association with allergic inflammation such as asthma in contemporary human populations^37^. According to the previous GWAS on eosinophil counts in the Japanese population, eosinophil counts had a significant genetic correlation with the asthma risk^28^. The relationship of mean *JAS* values with the obesity rate among 5-year-old children for each prefecture (Annual Report of School Health Statistics Research for 2021 academic year, URL: https://www.e-stat.go.jp/stat-search/files?page=1&query=%E8%82%A5%E6%BA%80&layout=dataset&toukei=00400002&tstat=000001011648&stat_infid=000032216703&metadata=1&data=1) was presented in Figure 5D, and with the incidence rate of asthma exacerbations^38^ in Figure 5E. We found the significant correlations between *JAS* and above indices (*R* = 0.65, two-sided *t*-test *P* = 1.3 × 10^−6^ for obesity rate; *R* = −0.41, two-sided *t*-test *P* = 0.005 for incidence rate of asthma exacerbations), indicating that these regional differences in the modern Japanese phenotype are largely determined by differences in the Jomon ancestry proportions.

## Discussion

We developed the *AMI* as a summary statistic to detect the Jomon-derived variants in modern Japanese without requiring any genomic sequences from the former. The computer simulation showed that *AMI* can detect ancestral variants with high accuracy, even in an admixed population whose source populations diverged tens of thousands of years ago. We were also able to detect Jomon-derived variants by the *AMI* even changing the population history in the simulations. The evolutionary history of the Japanese archipelago population was somewhat controversial^5–7,9–11,13,39–41^, but whatever population history was correct, the present approach using the *AMI* would be possible to detect Jomon-derived SNPs. Moreover, it would also be able to apply to other admixed populations whose source populations diverged relatively recently. The genetic diversity of modern humans has been greatly influenced by population admixture events^31,42–45^. The *AMI* will be a powerful tool for the population history of not only the Japanese but also other admixed populations. It should be noted that we determined the threshold of the *AMI* by the Youden index based on coalescent simulations in present study, but one may set the threshold according to one’s own research purpose; if one wants to reduce false positives, one can set the threshold strictly; if one wants to reduce false negatives, one can set the threshold loosely. Practically, the *AMI* threshold does not necessarily have to be set based on simulations that assume a population history. In the regional comparison of modern Japanese, the threshold was set loosely to pick up as many Jomon-derived variants as possible in order to grasp the trend of the whole genome in each Japanese prefectural population. On the other hand, in allele frequency estimation of the Jomon people by the Jomon-derived haplotypes of modern Japanese, the threshold was set strictly because false positives of Jomon-derived variants may lead to incorrect estimation of Jomon-derived haplotype frequencies.

We proposed an allele frequency estimation method in ancestral populations by classifying haplotypes of the current population based on their origin. Combining allele frequencies from THC Jomon and continental ancestries with previous GWAS results, we reported several traits for which the Japanese ancestral populations were presumed to have exhibited a characteristic phenotype: height, triglyceride, blood sugar, CRP and eosinophil count. Regarding height, several morphological analyses suggested the short stature of the Jomon people^33–35^, and Isshiki et al. found that among regional populations in the Japanese archipelago, those genetically closely related to the Han Chinese had greater PS for height^16^. These previous studies gave a strong assurance of our finding that the Jomon people were genetically shorter than the continental ancestry. As for triglyceride and blood sugar, we inferred genetically higher triglyceride and blood sugar for the Jomon ancestry compared to the continental ancestry. The diet of the Japanese archipelago population seemed to have changed significantly to become more dependent on agricultural products with the introduction of rice cultivation by continental East Asians after the Late Jomon period^46–49^. In the Yayoi period, carbohydrate intake from crops increased, which affected the Japanese archipelago population in the Yayoi period, including a higher incidence of dental caries^50,51^. Based on these previous studies, the genetic characteristics of the Jomon people regarding their nutritional status could be successfully explained as follows, with reference to “the thrifty gene hypothesis”^52^; the Jomon ancestry of modern Japanese may have had more difficulty maintaining triglyceride and blood sugar levels with food foraging than continental East Asian ancestry with rice farming, and thus genetic factors for higher triglyceride and blood sugar levels would have been helpful for them. Concerning CRP and eosinophil counts, we found the continental ancestry to have genetically higher CRP and eosinophil counts than the Jomon ancestry. CRP is a pattern recognition molecule who has an important role to defend against bacterial infections^53,54^. Eosinophils are a variety of white blood cells and one of the immune system components that play an important role in the response to helminth infection^55^. In general, farming led to higher population densities, sedentarization, contact with neighboring populations and reduced out-of-camp mobility, resulting in the spread of virulent bacteria and helminths and greater exposure to these pathogens^56–58^. We speculate that the continental East Asian ancestry of modern Japanese, having begun rice farming prior to their migration to the Japanese archipelago, needed to increase their resistance to pathogens such as bacteria and helminth compared to the Jomon ancestry, and acquired genetic factors for increasing their CRP and eosinophil counts. This view is supported by some archaeological and evolutionary studies. The earliest skeletal tuberculosis in the Japanese archipelago was confirmed at the Aoyakamijichi site in the late Yayoi period (approximately 2,000 years ago)^59^. Suzuki and Inoue discussed that continental immigrants in the Japanese archipelago spread tuberculosis, describing that “Primary tuberculosis & produced serious damage to the prehistoric Jomon people and resulted in a rapid reduction of their indigenous population.”^59^. Concerning the helminth infection, the phylogenetic analysis of the mitochondrial DNA of the blood fluke *Schistosoma japonicum* indicated the dispersal of *Schistosoma japonicum* radiated from the middle and lower reaches of the Yangtze River, where rice farming originated, to various parts of East Asia occurred after 10,000 years ago with the spread of rice farming culture^60^. Kanehara and Kanehara found no roundworm eggs in soil samples at Sannai-Maruyama site despite the detection of many whipworm eggs^61^, so Matsui et al. argued that the prevalence of roundworm infection occurred after the Yayoi Period with the beginning of rice agriculture^62^. Following these, our results are quite convincing, considering as that continental immigrants were genetically adapted to their agricultural livelihood through polygenic selection to alleles that exhibit resistance to infectious diseases compared to the Jomon hunter-gatherers. We also found that differences in ancestry proportions have a significant influence on regional variation of obesity and asthma exacerbation in modern Japan. Combining these regional phenotype variations and the triglyceride, blood sugar and eosinophil counts, which showed significant difference between Jomon and continental ancestries, possible scenarios are as follows. For Jomon hunter-gatherers, increased triglyceride and blood sugar were important as a resistance to starvation, while for continental East Asian farmers increased CRP and eosinophil counts were important as a protection against infectious diseases. Continental East Asian farmers migrated into the Japanese archipelago during the Late Jomon and Yayoi periods with their rice-farming techniques, resulting in large-scale interbreeding with the indigenous Jomon hunter-gatherers. The regional populations of the current Japanese archipelago with a high ancestry proportion of Jomon have retained genetic factors for higher triglyceride and blood sugar, which resulted in a higher risk of obesity. On the other hand, the regional populations with a high ancestry proportion of continental East Asians have genetic factors that increase eosinophil counts, resulting in a higher risk of asthma exacerbation. Regional gradations in the obesity rate and incidence rate of asthma exacerbation among modern Japanese were caused by regional differences in the ancestry proportion of the continental East Asians. Overall, according to our findings with genome-wide allele frequencies and mean 2*βf* of the Jomon and continental ancestries, we can conclude that (1) some phenotypes of Jomon people and continental East Asians were highly diverged at the genome-wide scale (2) some phenotypic differences may have been the result of genetic adaptations to the respective livelihoods of the Jomon people and continental East Asians (3) regional variations in admixture proportions of the Jomon people and continental East Asians formed phenotypic gradations of current Japanese archipelago populations. Integrating future GWAS of other traits in modern Japanese with allele frequencies on the Jomon ancestry would reveal their phenotypic characteristics that do not appear in excavated skeletal morphology. It is also possible, perhaps, to discover further regional gradations in phenotypes of modern Japanese caused by regional differences in admixture proportions, as in the case of height, obesity rate and asthma exacerbation.

As for the process of population formation in the Japanese archipelago from the Late Jomon period to the present, we propose a model, which is shown in Figure 6. From the Late to Final Jomon period, the Jomon hunter-gatherers settled down in mainland Japan. They were a relatively short statured population with genetic factors to adapt to their hunter-gatherer lifestyle, such as higher triglyceride and blood sugar for resistance to starvation. The population size or the population density of the Jomon people varied among regions, relatively large in Tohoku and Kyushu and relatively small in Kinki, Shikoku^23^. At the same time, rice-farming populations lived in continental East Asia; they had relatively tall stature and genetically adapted to their livelihood with higher CRP and eosinophil counts for protecting against helminths. In the Final Jomon period, the continental East Asians arrived in northern Kyushu with their rice-farming technique and then admixed with the Jomon people in all the regions of mainland Japan. During the Yayoi period, the population size of immigrants was relatively increased in the Kinki and Shikoku regions, where the populations were small at the end of the Jomon period. In the Kinki and Shikoku regions, rice farming seemed to have started relatively early compared to other regions^25^. Regional differences in population size from the Final Jomon to the Yayoi period made a variation in admixture proportions of the Jomon people and continental East Asians among regions in modern Japanese archipelago. Regional variations in admixture proportion have resulted in the geographic gradations of today’s Japanese genotypes and phenotypes.

**Figure 6.**
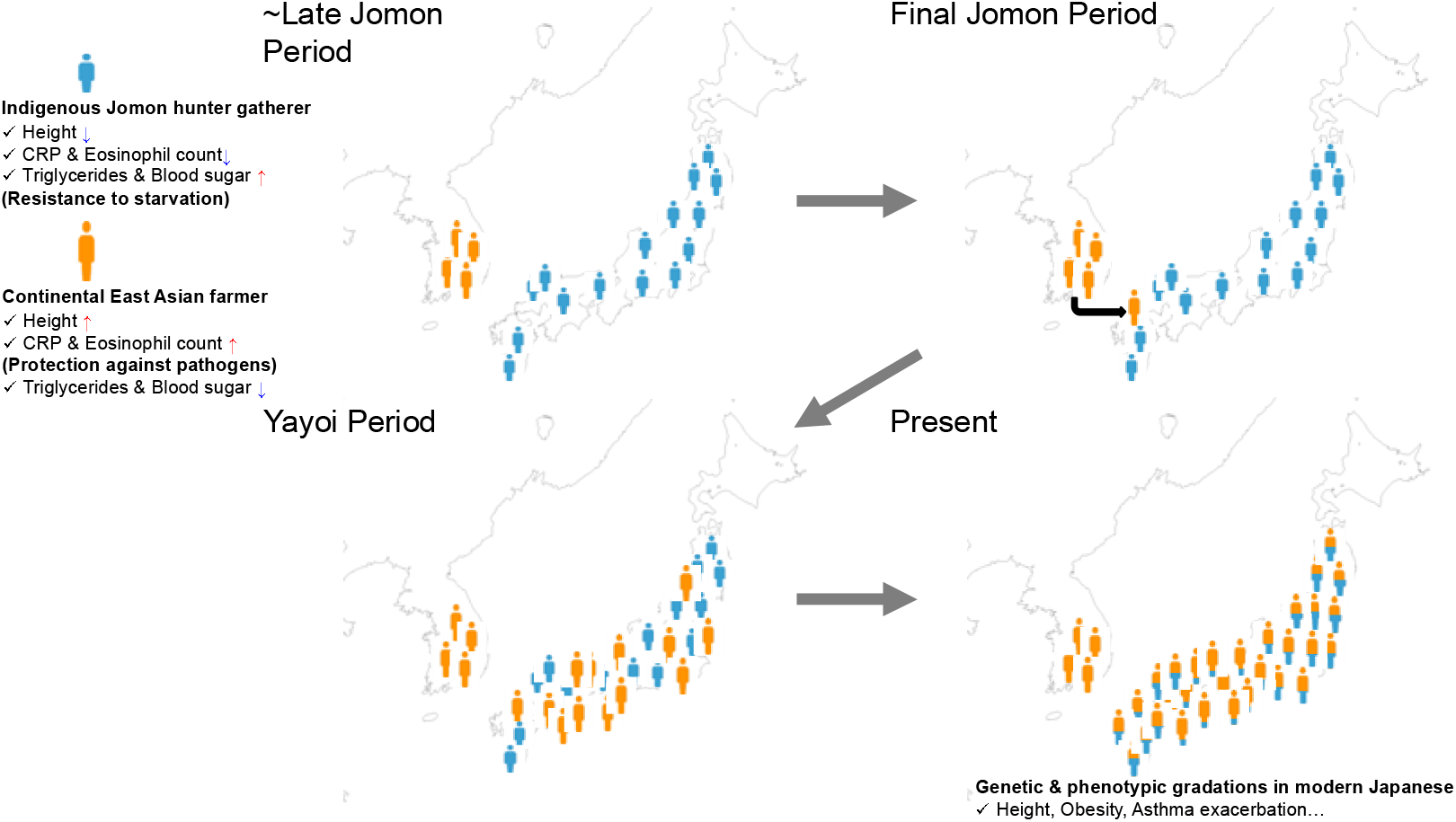
Formation process of regional population of the mainland Japan. (Upper left panel) In the Jomon period, the Jomon hunter-gatherers, who were a relatively short statured population with genetic factors such as high blood sugar, settled down in mainland Japan, while rice-farming populations, who had relatively tall stature and genetic factors such as higher eosinophil counts, lived in continental East Asia. These genetic factors seemed to be adaptive to their respective lifestyles. (Upper right panel) In the Final Jomon period, the continental East Asians arrived in northern Kyushu and admixed with the Jomon people in mainland Japan. (Lower left panel) During the Yayoi period, the population size of immigrants was relatively increased in the Kinki and Shikoku regions, where the populations were small at the end of the Jomon period. (Lower right panel) Regional variations in admixture proportion, which may be caused by regional differences in population size from the Final Jomon to the Yayoi period, have resulted in today’s geographic gradations of mainland Japanese genotypes and phenotypes.

## Author Contributions

Y.W. and J.O. conceived the study. Y.W. designed and conducted the data analyses. Y.W. performed the computer simulations. Y.W. wrote the manuscript with support from J.O. J.O. supervised the project. All authors read and approved the final manuscript.

## Acknowledgments

We are grateful to the individuals who participated in the study. We would like to express our deepest gratitude to Mr. Masahiro Inoue, Shota Arichi, and Akito Tabira who obtained the genotype data and provided the technical environment for analyzing them. We would like to thank Dr. Hiroki Ota of Tokyo University, Tokyo, Japan, and Dr. Takashi Gakuhari of Kanazawa University, Ishikawa, Japan, for providing us the BAM file of the IKawazu Jomon. We also thank Dr. Naruya Saito from the National Institute of Genetics, Shizuoka, Japan, and Dr. Hideaki Kanzawa-Kiriyama from the National Museum of Nature and Science, Tokyo, Japan for providing us the BAM file of the Funadomari Jomon. We also thank Dr. Nakagome Shigeki from the the School of Medicine, Trinity College Dublin, Dublin, Ireland for providing us the BAM file of the nine Jomon and three Kofun individuals. Computations were partially performed on the NIG supercomputer at ROIS National Institute of Genetics. This study was partly supported by Grant-in-Aid for Scientific Research (B) (18H02514) and Grant-in-Aid for Scientific Research on Innovative Areas (19H05341 and 21H00336) and Grant-in-Aid for Early-Career Scientists (21K15175) from the Ministry of Education, Culture, Sports, Science, and Technology of Japan.

## Materials and Methods

### Samples

All the individuals investigated in this study were customers of the Japanese Direct to Consumer (DTC) genetic-testing service, HealthData Lab (Yahoo! Japan Corporation, Tokyo, Japan). They were provided an agreement, and informed consent was obtained for their data to be used for research. In this study, the Japanese archipelago was divided into eleven regions (Figure S1 and Table S1): Hokkaido (430 individuals), Tohoku (746 individuals), Kanto (3,990 individuals), Hokuriku (431 individuals), Chubu (410 individuals), Tokai (933 individuals), Kinki (1,861 individuals), Chugoku (600 individuals), Shikoku (314 individuals), Kyushu (1,016 individuals), and Okinawa (111 individuals). All statistical analyses were conducted at the Yahoo! Japan Corporation, with personal information of the customers completely hidden. We obtained approval from the Ethics Committee of the Yahoo! Japan Corporation. The individual genotypes of 10,842 Japanese analyzed in this study are not available to avoid personal identification.

### Data processing

In this study, we used genotype information from about Japanese individuals who had already been genotyped in a previous study^15^. This section briefly outlines the genotype data processing (see Watanabe et al. 2021 for more detailed description). For the 11,069 Japanese in total, saliva samples were collected using Oragene®-DNA (OG-500) (DNA Genotek, Ottawa, Canada) and DNA was extracted in accordance with the manufacturer’s instructions. The Illumina HumanCore-12 Custom BeadChip and HumanCore-24 Custom BeadChip (Illumina, San Diego, CA), were used for the genotyping. The genotype data was filtered by PLINK version 1.9 at Hardy-Weinberg equilibrium *P* value□<□0.01, SNP call rate□<□0.01, and sample call rate□<□0.1 (QC phase 1). Further, 116 samples that were genetically close to Han Chinese in the PCA and 111 samples with higher IBD values than 0.0125 with one or more subjects were excluded, and 10,842 samples were used for the subsequent analyses.

### Coalescent simulation by msprime

To investigate the characteristics of the Jomon-derived autosomal genomic components of mainland Japanese, we conducted a coalescent simulation assuming the admixture of the Jomon and continental East Asians using msprime^63^ (Figure S2). A remarkable feature of the msprime program is that it specifies the time and population where the mutation and coalescence events occurred. Our simulation code was made with reference to a previous study^64^. Our custom code for msprime simulation was described in Supplementary Text. The split between the Jomon ancestors and continental East Asians was set to 1,200 generations ago (30,000 YBP), according to the divergence time (between 18,000 YBP and 38,000 YBP) estimated in Kanzawa-Kiriyama et al.^10^ and the beginning of the Jomon period (around 16,000 YBP) ^2^. Migration from continental East Asia to mainland Japan was set between 120 and 80 generations ago, with reference to the beginning of the Yayoi period, around 2,800 years ago ^4^. The total admixture proportion of the Jomon people in the modern mainland Japanese was set to 12%^8^. The effective population size was set to 5,000 for both populations. The mutation rate and recombination rate were set to 1.2×10^−8^ per bp per generation and 1.3×10^−8^ per bp per generation, respectively^65–68^.

This study aimed to detect Jomon-derived variants based on LD among Japanese specific variants. There are three types of Japanese specific variants: (type 1) Jomon-derived variants; (type 2) variants derived from continental East Asians; and (type 3) novel variants (Figure 1). It should be noted that Japanese specific variants generated earlier than the split time of the Jomon people and the continental East Asians were classified as Jomon-derived variants (type 1). We compared the LD status of three types of Japanese specific variants in the coalescent simulations. The origin of each haplotype of mainland Japanese can be estimated from coalescent time to the haplotypes of the Jomon or continental East Asians. That is, if a haplotype of a mainland Japanese sample coalesced with haplotypes of Jomon samples earlier than the admixture of the Jomon people and continental East Asians, the haplotype is inferred to be derived from Jomon. To extract the three types of Japanese specific variants (i.e., variants not found in samples from continental East Asians), 3,000 replicates of 1 Mb simulations were performed. We sampled 200 haplotypes from each of the four populations (modern mainland Japanese, modern continental East Asians, Jomon people 120 generations ago, and continental East Asians 120 generations ago^15^) to detect variants observed in modern mainland Japanese but not seen in continental East Asians. Each Japanese specific variant was classified into (type 1) the Jomon-derived variant, (type 2) the continental East Asian-derived variant, and (type 3) the novel variant based on when and in which lineage the mutation occurred (Figure 1). To calculate the ancestry marker index (*AMI*), we first calculate the LD coefficient *r*^2^ between Japanese specific variant pairs within each 1 Mb bin. For each type of the Japanese specific variant, the *AMI* was calculated as:

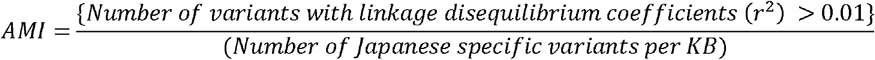

Jomon-derived variants are expected to have higher *AMI* values. The performance of the *AMI* was verified by receiver operating characteristic (ROC) analysis using the ROCR package in R. The threshold to detect Jomon-derived variants was determined based on the Youden Index.

### Detection of Jomon-derived variants in real data

Jomon-derived variants were inferred from the whole genome sequence data from 26 populations from different parts of the world, including mainland Japanese (JPT) and four continental East Asian populations (CHB, CHS, CDX, and KHV), obtained from 1KG Phase III^22^, and 87 individuals from the KPGP^69^. In this study, only biallelic SNPs were used. Prior to extracting the Jomon-derived variants, we performed a PCA in PLINK (version 1.9)^70^ using 1KG mainland Japanese (JPT) and Han Chinese (CHB) data. During this analysis, we found that one JPT individual (NA18976) was close to the continental East Asians (Figure S8), so NA18976 was excluded from subsequent analyses. First, 1,784,634 SNPs specific to 1KG JPT were detected using VCFtools v0.1.13^71^. Next, LD coefficients (*r*^2^) were calculated between the Japanese specific SNPs located within 1 Mb from each other with the --hap-r2 option of VCFtools in combination with the --ld-window-bp option. The number of SNPs with *r*^2^ > 0.01 was counted for each Japanese specific SNP. The density of Japanese specific variants per 1 kb of each chromosome was calculated using the --SNPdensity option of VCFtools, and the *AMI* was calculated for each Japanese specific SNP. To eliminate the possibility of sequence errors, regions with a density of Japanese specific variants per kb below a mean of - 1sd of each chromosome were excluded from the analysis. In this analysis, we assumed that the number of Japanese specific variants per kb, which is the denominator of the *AMI*, is constant for each chromosome (i.e., the numerator of the *AMI* was normalized for each chromosome). Based on the threshold set by the ROC analysis of simulated Japanese specific variants, we inferred SNPs originating from the Jomon people in the real data.

### Verification of Jomon-derived variants based on whole-genome sequence data from Jomon remains

For the verification of Jomon-derived variants based on the whole genome sequence data, we devised the “Jomon allele score” (*JAS*). The *JAS* was calculated using the following formula:

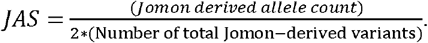

We calculated *JAS* for the Ikawazu^9,11^ and Funadomari^10^ Jomon, as well as for 104 individuals from the 1KG JPT. The BAM file of the Ikawazu Jomon was provided by Hiroki Ota of Tokyo University, Tokyo, Japan, and Takashi Gakuhari of Kanazawa University, Ishikawa, Japan. The BAM file of the Funadomari Jomon was provided by Naruya Saito from the National Institute of Genetics, Shizuoka, Japan, and Hideaki Kanzawa-Kiriyama from the National Museum of Nature and Science, Tokyo, Japan. The genotypes of Ikawazu Jomon and Funadomari Jomon samples were called by the UnifiedGenotyper tool in the GenomeAnalysisToolkit version 3.6^72^. For the Ikawazu Jomon, the --mbq 30 --ploidy 2 --output_mode EMIT_ALL_CONFIDENT_SITES options were specified. For the Funadomari Jomon, the options described in the original paper were specified. Jomon-derived variants were subjected to LD pruning by the --indep-pairwise command of PLINK (--indep-pairwise 1000 200 0.8). In addition, only the Jomon-derived variants of depth ≥ 6 in the Ikawazu and Funadomari Jomon were used for the calculation of the *JAS*. As a result, 4,458 SNPs were used to calculate *JAS*.

### Imputation of genotypes of Jomon-derived variants

Haplotype phasing and genotype imputation were performed using EAGLE2^73^ and Minimac3^74^, respectively, with whole genome sequence data of 413 mainland Japanese^26^ phased by SHAPEIT2^75^. After the imputation, Jomon-derived variants with high imputation quality (R^2^ > 0.8) were extracted. Also, LD pruning was performed with PLINK (--indep-pairwise 1000 200 0.1), and a total of 3,917 Jomon-derived variants were used for the analysis.

### Geographical distribution of the *JAS*

In subsequent analyses, individuals from Hokkaido that were largely affected by immigration after the Meiji period were excluded. Using 3,917 Jomon-derived variants, we calculated the *JAS* for individuals of each prefecture and compared them between regions and prefectures.

We compared the population size estimated from the number of archeological sites in each prefecture, assuming that the population size per archeological site was constant in each prefecture during the same period. We examined the correlations between (A) the average *JAS* in each prefecture and the number of archeological sites from the Jomon period (obtained from the Statistical report of buried cultural properties, Agency of Cultural Affairs, Japan; http://www.bunka.go.jp/seisaku/bunkazai/shokai/pdf/h29_03_maizotokei.pdf), (B) the average *JAS* in each region and the population size estimated from the number of archeological sites in the Late Jomon period^23^, and (C) the average *JAS* in each prefecture and the log_10_(number of archeological sites in the Yayoi period/number of archeological sites in the Late Jomon period)^23^. Finally, (A) and (C) were plotted for each prefecture, while (B) was plotted for each region because data for each prefecture were not available. Correlation test was conducted by R cor.test function (df = 43). We further compared the average *JAS* and the arrival date of rice farming in each mainland Japanese region which was estimated based on the radiocarbon dating of charred rice remains in Crema et al.^25^. In this comparison, we adopt the regional classification of Crema et al. rather than the regional classification used in other analyses of this study.

### Inference of phenotypic characteristics of the Jomon and continental East Asian ancestries by estimating their allele frequencies of genome-wide SNPs by Jomon-derived haplotypes

First, we applied the *AMI* to detect Jomon-derived variants in the whole genomes of 419 modern Japanese (THC dataset, *AMI* threshold = 100). The haplotypes of the regions around a focal SNP site (circles in Figure S13) were classified as “Jomon-derived haplotypes” or “continental haplotypes”, depending on the presence of Jomon-derived variants in the 10 kb region upstream or downstream of the focal SNP. We extracted 67,607 Jomon-derived variants for estimating allele frequencies in THC Jomon ancestry and THC continental ancestry. We inferred allele frequencies of 6,481,773 genome-wide SNPs of minor allele frequencies > 1% in modern Japanese. The allele frequency estimation was performed by R.

The estimated allele frequencies in the THC ancestries were compared with those in the 11 Jomon^9–11,13^, three Kofun^13^, THC 419 modern Japanese^26^ and 103 Han Chinese (CHB from 1000 Genome Project) individuals^22^. In addition to the already genotyped Ikawazu and Funadomari Jomon individuals, bam files of nine Jomon and three Kofun individuals were provided by Dr. Nakagome of the School of Medicine, Trinity College Dublin, Dublin, Ireland, and the genotypes were determined by GATK UnifiedGenotyper. In the pairwise f3(A, B, Yoruba (= YRI in 1KG)) test, we used 32,143 SNPs that were genotyped in all the individuals of Jomon, Kofun and modern individuals.

To estimate the phenotypic characteristics of the Japanese ancestral population, we combined genome-wide allele frequencies from THC Jomon and continental ancestries with GWAS for 60 quantitative traits in modern Japanese population(Table S1)^27–29^. First, referring to the QTL GWAS results for Japanese from previous studies, LD-based clumping was performed using the --clump option in PLINK to extract phenotype-associated SNPs with no LD in modern Japanese of 1000 Genome Project^22^ for each trait, setting *P* value threshold as 0.01, 0.001 and 0.0001. We then calculated following index using *n* SNPs with lower *P* value than threshold we set (P < 0.01, *P* < 0.001 and *P* < 0.0001) for each trait:

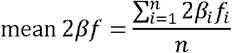

Here *β_i_* represents the effect of the *i*-th SNP of previous QTL GWAS results, and *f_i_* represents the effect allele frequency for the *i*-th SNP. The mean 2*βf* represents the average phenotype of a focal population, and the phenotype of each population can be compared by the magnitude of this value. For each trait, we calculated the mean 2*βf* values for THC modern Japanese, Jomon and continental ancestries. The next step was to identify traits with significantly large differences in mean 2*βf* between THC Jomon and continuous ancestries. For each trait, the null distribution of mean 2*βf* was estimated by 1,000 simulations, where the allele frequency *f_i_* for each SNP was randomly switched to THC Jomon or continental ancestries and calculated mean 2*βf* values. Based on the 97.5th and 2.5th percentile of the null distribution of mean 2*βf*, the following index *D* was calculated for 60 traits to verify the magnitude of phenotypic differences between THC Jomon and continental ancestries:

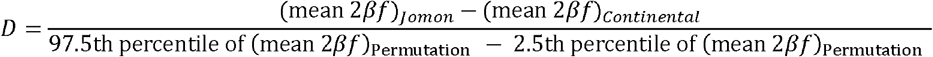

The larger the absolute value of *D*, the more different between Jomon and continental ancestries is for a focal trait.

## Data and code availability

The individual genotypes of 10,842 Japanese analyzed in this study are not available to avoid personal identification. The list of Jomon-derived variants detected in this study, and the allele frequencies of Jomon-derived variants in each Japanese prefecture are available from the lead contact upon request. Our custom code for msprime simulation was described in supplemental information. Any additional information required to reanalyze the data reported in this paper is available from the lead contact upon request.

