## Supplemental information for "Modern Japanese ancestry-derived variants revealed the formation process of the current Japanese regional gradations"

**Confirmation of the performance of S* in mainland Japanese population by coalescent simulation**

S* is a summary statistic used to detect genomic segments of anatomically modern humans which derived from archaic hominins with a divergence time about 1 million years ago^1,2^. We confirmed the performance of S* in the admixture of two relatively recently divergent populations, the Jomon people and the continental East Asians, by msprime coalescent simulations^3^ assuming the Japanese population history (Figure S2). Two patterns of divergence time between the Jomon people and the continental East Asians was compared in the coalescent simulations; (a) 40,000 generations ago (1,000,000 YBP, assumed the divergence of archaic hominin and anatomically modern humans) and (b) 1,200 generations ago (30,000 YBP, assumed the divergence of the Jomon people and the continental East Asians). Ten independent replicates of 1 Mb simulation were conducted (i.e., chromosomes 1 Mb in length were simulated 10 times) in each pattern of the divergence time. Output vcf files containing 1 Mb genotypes of the mainland Japanese and the continental East Asians were divided into 50 kb windows, and S* was calculated for each generated mainland Japanese sample. Coalescent simulations implemented by using mspime recorded whether the generated mainland Japanese haplotypes are derived from the Jomon people or the continental East Asians. For each simulation with the two patterns of divergence times, we compared S* between samples with homozygote of continental East Asian-derived haplotypes, samples with heterozygote of continental East Asian-derived haplotypes and Jomon-derived haplotypes, and samples with homozygote of Jomon-derived haplotypes (Figure S3). In pattern (a), the distributions of S* were extremely different between samples of homozygote of East Asian-derived haplotypes and other samples (Figure S3A), while the distributions of S*were very similar between the three samples in pattern (b) (Figure S3B). These results suggested that in the case of populations derived from the admixture of two relatively recently divergent populations (i.e. the admixture between anatomically modern humans), it is not possible to distinguish genomic segments derived from admixture by S*. S* assumes admixture between an archaic hominin and *Homo sapiens* with a divergence time of approximately 500 thousand to 1 million years ago, so it seems that an insufficient number of Jomon-derived specific variants of mainland Japanese which accumulated in the Jomon-lineage causes reduction in power to detection.

Based on the principle of S*^1^, we assumed that, to detect ancestry-derived segments with S*, there should be a large number of ancestry-derived variants on the ancestry-derived segments as genetic markers. (Here, "ancestry-derived" refers to “archaic-hominin-derived” in the case of admixture between archaic hominins and modern humans, and “Jomon-derived” in the case of admixture between the Jomon people and continental East Asians.) Comparing (1) admixture between archaic hominins and modern humans, which is the subject of S*, and (2) admixture between the Jomon people and continental East Asians, which is the subject of our study, archaic hominins and modern humans (target populations in (1)) diverged much earlier (500,000 years ago~) and much more lineage-specific mutations have accumulated in archaic hominins than in case of (2), so the number of ancestry-derived variants per ancestry-derived segment is expected to be higher in (1) than in (2). We ran additional simulations to examine the relationship between the number of ancestry-derived SNPs per segment and the value of S*. First, assuming a population history of (1) and (2), we ran 1 Mb coalescent simulations ten times, respectively. Here, in (1), the divergence time of archaic hominins and modern humans was set to 40,000 generations ago, and the ancestry proportion of archaic hominins in modern humans is set to 4%. Parameters in (2) are the same as Figure S2. Next, we divided the generated sequences into 50 kb windows. Of each 50 kb window, we extracted only the window containing the “true” ancestry-derived segment, and counted the number of “true” ancestry-derived variants in the window, and calculated S* value for each individual. Since segments including individuals with higher S* value are detected as ancestry-derived segments, we focused on the maximum value of S* in each 50 kb window (S*max) and plotted the number of ancestry-derived variants and S*max in each window (Figure S4). This result indicates that at least a certain number of ancestry-derived variants per segment must be present for the larger S* value and that the number of ancestry-derived variants per segment is smaller in (2) than in (1). Based on this result, we can conclude as follows; in the case of admixture between archaic hominins and modern humans, there are many archaic-Hominin-derived variants per segment, which can detect archaic-Hominin-derived segments by S*; in the case of admixture between the Jomon people and continental East Asians, the number of Jomon-derived variants per segment is small, which cannot detect Jomon-derived segments by S*.

**Geographic distribution of *JAS in* higher *AMI* cutoff value**

By increasing the cutoff value of *AMI*, more robust Jomon-derived variants can be detected. When the cutoff value was changed to 100, 474 Jomon-derived variants were detected in 10,412 samples. The *JAS* values were recalculated for these SNPs and compared with those from the *AMI* cutoff value of 28.0374. No significant change was observed in the order of the *JAS* values by prefecture (Figure S10), suggesting that the geographical distribution of the *JAS* does not depend on the AMI cutoff.

**Python code of 1 Mb simulation of the Japanese population history used for confirmation of Jomon derived haplotype length distribution in this study**

*# python Jomon_coales_sim.py seed outprefix*

\# Confirmed working with python 2.7.5

import sys,msprime,gzip

from math import exp

from collections import OrderedDict

import numpy as np

import itertools

nhy = 200 *# num. sampled Continental East Asian haplotypes in 120 generations ago*

nhja = 200 *# num. sampled Japanese haplotypes*

nhjo = 200 *# num. sampled Jomon haplotypes in 120 generations ago*

nhko = 200 *# num. sampled Continental East Asian haplotypes*

nbp = 1000000 *# simulated 1 Mb*

rho = 1.3e-8 *# recombination rate*

mu = 1.2e-8 *# mutation rate of base simulation*

seed = sys.argv[1]*# we used seeds 1-3000*

outprefix = sys.argv[2] + "_" + sys.argv[1]*# output file prefix*

NY = 5000 *# Continental East Asian haplotypes effective size*

NJ = 5000 *# Jomon eff. size*

*# times are in generations before present*

TJY = 1200 *# Jomon-Continental East Asian split time*

Tadmixstart = 120 *# start of introgression*

Tadmixend = Tadmixstart-40 *# end introgression*

*##Tgrowth = 200 # time of recent growth*

*# migration rates are proportion of population made of new immigrants each generation*

mintro = 0.051626236 *# introgression from continental East Asians into Jomon*

pop_config = [ msprime.PopulationConfiguration(sample_size = None, initial_size = NJ), msprime.PopulationConfiguration(sample_size = None, initial_size = NY)]

samples = [msprime.Sample(0,0) for i in range(nhja)] + [msprime.Sample(1,0) for i in range(nhko)] + [msprime.Sample(0,Tadmixstart) for i in range(nhjo)] + [msprime.Sample(1,Tadmixstart) for i in range(nhy)] *# Jomon-Continental East Asian haplotypes sampled 120 gen. ago*

*# Continental East Asian haplotypes introgression*

admix_event = [msprime.MigrationRateChange(time = Tadmixend, rate = mintro, matrix_index = (0,1)), msprime.MigrationRateChange(time = Tadmixstart, rate = 0, matrix_index = (0,1))]

*# Jomon Continental East Asian haplotypes split*

JY_event = [msprime.MassMigration(time=TJY, source=1, destination=0)]

events = admix_event + JY_event

*# make the simulation*

treeseq = msprime.simulate(population_configurations = pop_config, samples = samples, demographic_events = events, length = nbp, recombination_rate = rho,mutation_rate = mu, random_seed = seed)

*#output the vcf*

with gzip.open(outprefix+".vcf.gz","w") as vcffile:

treeseq.write_vcf(vcffile,2)

jp=treeseq.samples()[0:nhja]

jpt=treeseq.simplify(samples=jp)

JSV=OrderedDict() *# Japanese specific variants*

JSV["jn"]=OrderedDict() *#jn, type 3*

JSV["jd"]=OrderedDict() *#jd, type 1, occurred after the split of Jomon and continental East Asians*

JSV["yd"]=OrderedDict() *#yd, type 2, occurred after the split of Jomon and continental East Asians*

JSV["jt"]=OrderedDict() *#jt, type 1, occurred before the split of Jomon and continental East Asians*

JSV["yt"]=OrderedDict() *#yt, type 2, occurred before the split of Jomon and continental East Asians*

mf=OrderedDict() *# frequency of Japanese specific variants*

mf["jn"]=OrderedDict()

mf["jd"]=OrderedDict()

mf["yd"]=OrderedDict()

mf["jt"]=OrderedDict()

mf["yt"]=OrderedDict()

*#extract Japanese specific variants*

for var in treeseq.variants():

n=var.site.mutations[0].node

t=treeseq.node(n).time

p=treeseq.node(n).population

gj=var.genotypes[0:200]

gk=var.genotypes[200:400]

gjp=var.genotypes[400:600]

if sum(gj)>0 and sum(gk)==0:

if t > Tadmixstart and t < TJY and p==0:

JSV["jd"][var.index] = var.site.position

mf["jd"][var.index]=float(sum(gj))/len(gj)

elif t > Tadmixstart and t < TJY and p==1:

JSV["yd"][var.index] = var.site.position

mf["yd"][var.index]=float(sum(gj))/len(gj)

elif t < Tadmixstart and p==0:

JSV["jn"][var.index] = var.site.position

mf["jn"][var.index]=float(sum(gj))/len(gj)

elif t > TJY and p==0 and sum(gjp)>0:

JSV["jt"][var.index] = var.site.position

mf["jt"][var.index]=float(sum(gj))/len(gj)

elif t > TJY and p==0 and sum(gjp)==0:

JSV["yt"][var.index] = var.site.position

mf["yt"][var.index]=float(sum(gj))/len(gj)

pos=OrderedDict()

pos["jd"]=OrderedDict()

pos["yd"]=OrderedDict()

pos["jn"]=OrderedDict()

pos["jt"]=OrderedDict()

pos["yt"]=OrderedDict()

*#record LD coeff*

for var in jpt.variants():

p=var.site.position

pos_jd=JSV["jd"].values()

pos_yd=JSV["yd"].values()

pos_jn=JSV["jn"].values()

pos_jt=JSV["jt"].values()

pos_yt=JSV["yt"].values()

if any (elem == p for elem in pos_jd):

pos["jd"][var.index] = p

elif any (elem == p for elem in pos_yd):

pos["yd"][var.index] = p

elif any (elem == p for elem in pos_jn):

pos["jn"][var.index] = p

elif any (elem == p for elem in pos_jt):

pos["jt"][var.index] = p

elif any (elem == p for elem in pos_yt):

pos["yt"][var.index] = p

for type in pos.keys():

l=len(pos[type])

l=range(0,l)

com=itertools.combinations(l,2)

wf=open(outprefix+"_r2.array."+type+".txt","w")

for c in com:

v0=pos[type].keys()[c[0]]

v1=pos[type].keys()[c[1]]

r2=str(msprime.LdCalculator(jpt).r2(v0,v1))

pos0<-str(pos[type].values()[c[0]]

pos1<-str(pos[type].values()[c[1]])

print >> wf,"%.1f"%pos0,"%.1f"%pos1,"%.1f"%r2,

wf.close()

com=itertools.combinations(pos.keys(),2)

for c in com:

wf=open(outprefix+"_r2.array."+c[0]+c[1]+".txt","w")

l0=len(pos[c[0]])

l0=range(0,l0)

l1=len(pos[c[1]])

l1=range(0,l1)

for i in l0:

for j in l1:

v0=pos[type].keys()[c[0]]

v1=pos[type].keys()[c[1]]

r2=str(msprime.LdCalculator(jpt).r2(v0,v1))

pos0<-str(pos[type].values()[c[0]]

pos1<-str(pos[type].values()[c[1]])

print >> wf,"%.1f"%pos0,"%.1f"%pos1,"%.1f"%r2,

wf.close()

**Supplemental Figures**

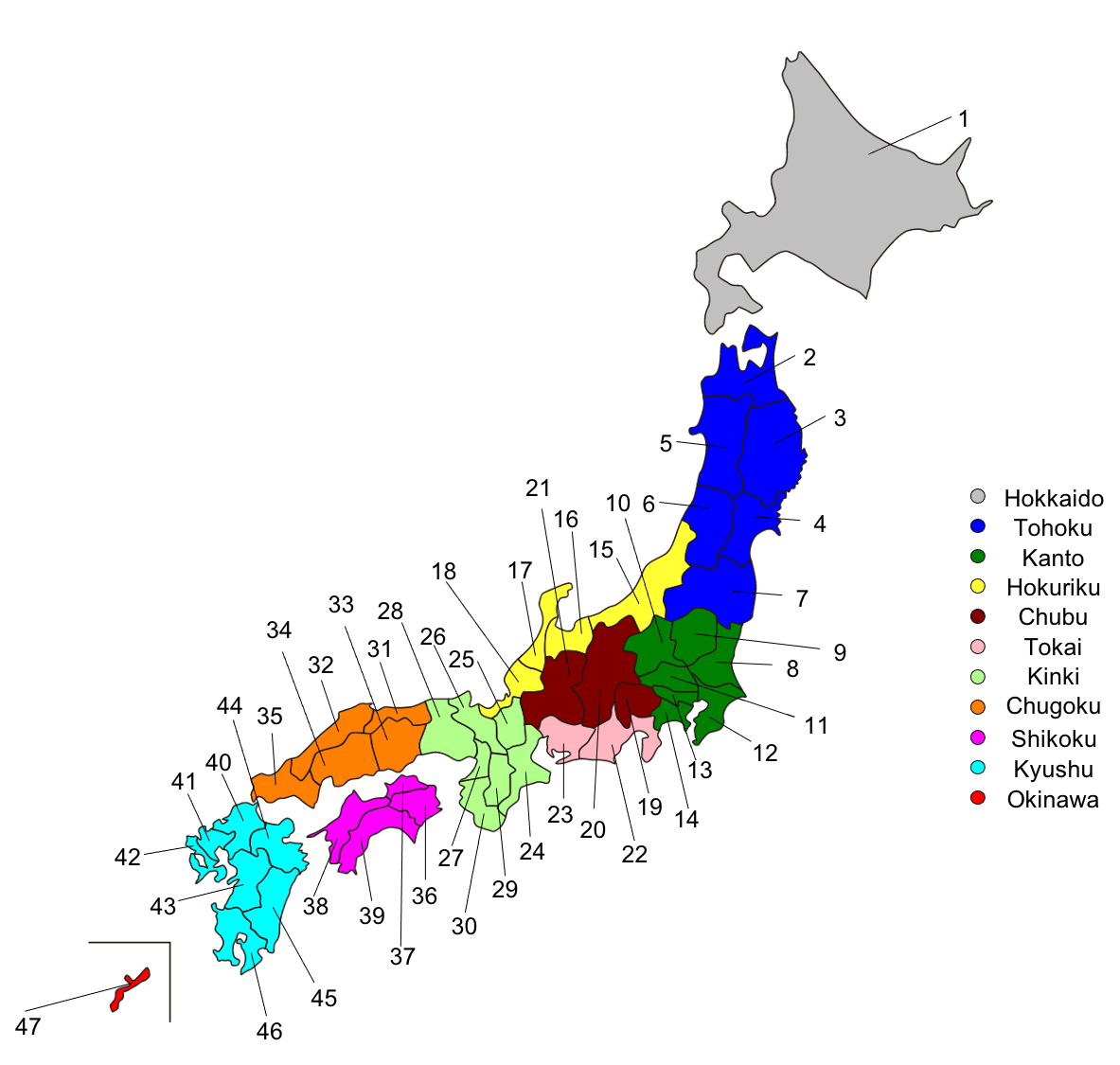

**Figure S1.** Map of the Japanese prefectures. The prefectures of Japan are divided into eleven regions. The prefecture numbers in Supplementary Table 1 are indicated (the corresponding prefecture names are given in Supplementary Table 1). In this study, “mainland Japanese” means the Japanese people except for individuals from Hokkaido and Okinawa.

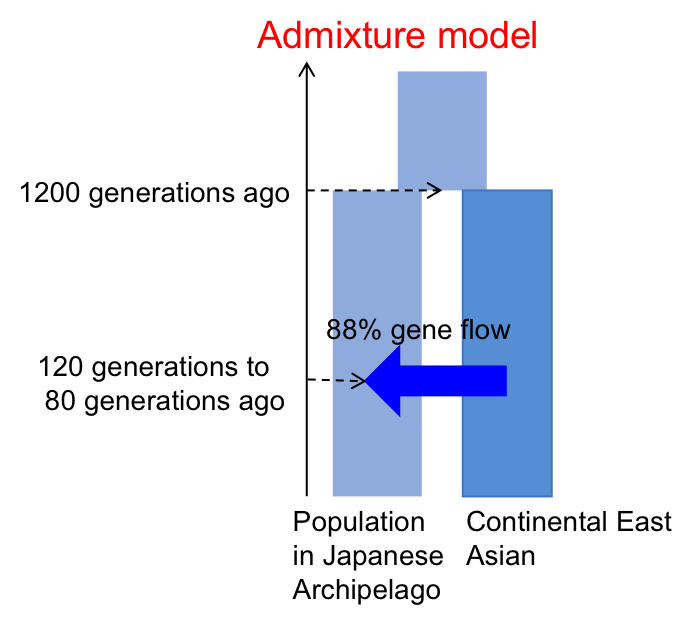

**Figure S2.** Basic demographic model of the Japanese population assumed in my coalescent simulations. The split between the Jomon ancestors and the continental East Asians was set to 1,200 generations ago (30,000 YBP). The continuous migration from the continental East Asian to the Jomon was set between 120 and 80 generations ago (3,000 ~2,000 YBP). The total admixture proportion of the Jomon people in the modern Mainland Japanese was set to 12%. The effective population size was 5,000 for each population. Some parameters were varied as necessary for the analysis of Figure S3, S4, and S6 to validate the performance of S* and the *AMI*.

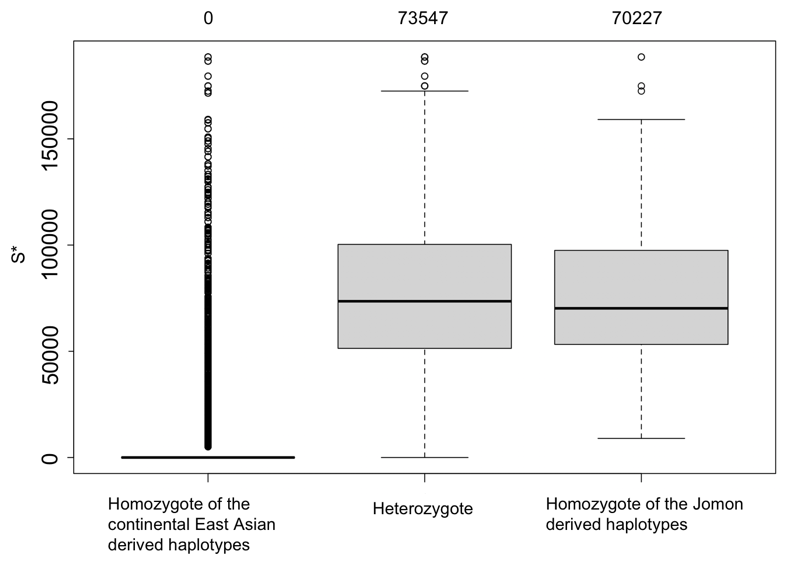

(A)

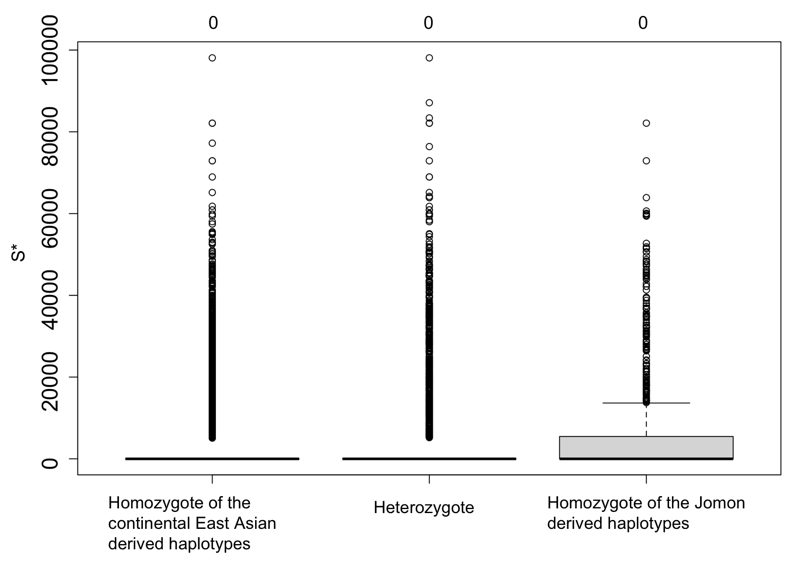

(B)

**Figure S3.** Distribution of S* in simulated data assumed two patterns of divergence time in each sample. The boxplots of the S* in homozygotes of the continental East Asian-derived haplotypes, the heterozygotes of the continental East Asian-derived haplotype and the Jomon-derived haplotypes, and the homozygotes of the Jomon-derived haplotypes is presented. (A) and (B) assumed divergence time 40,000 and 1,200 generations ago, respectively. The number on each boxplot shows the median of S* in each sample.

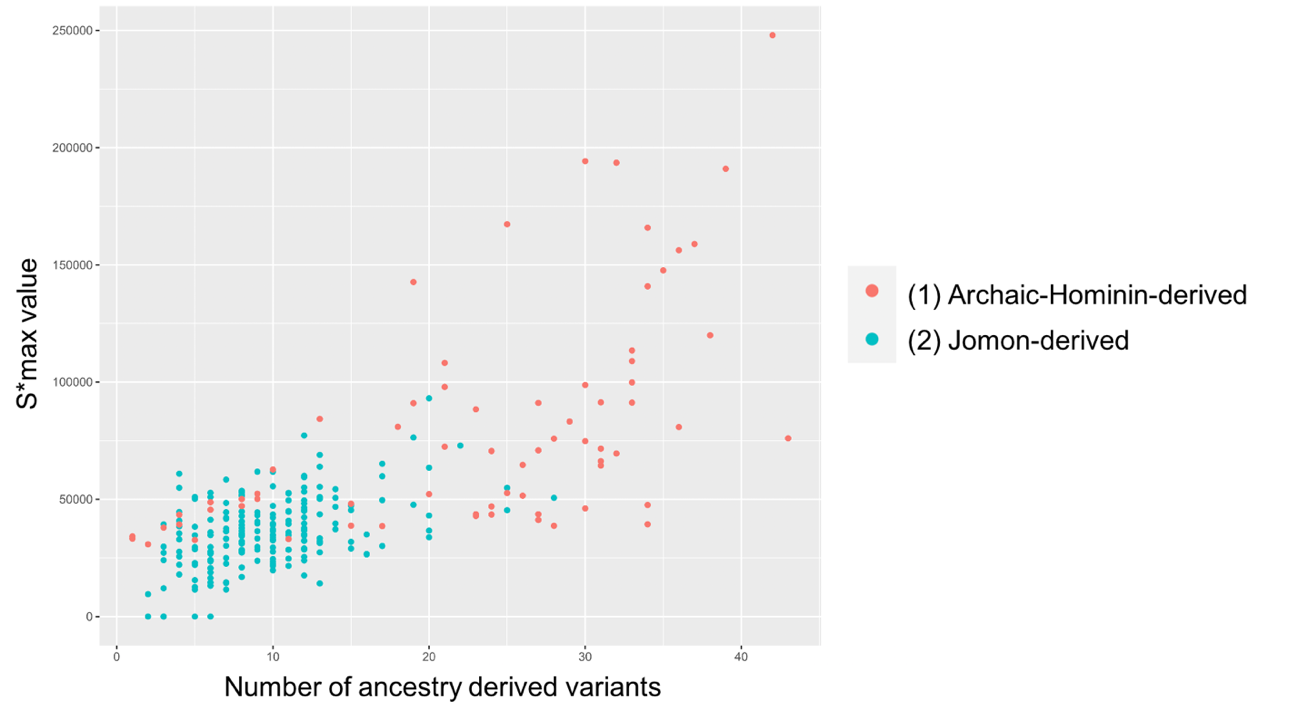

**Figure S4.** Relationship between the number of ancestry derived variants and the S* max value in each “true ancestry-derived” 50 kb segment in simulated data assumed two patterns of divergence time. (A) and (B) assumed divergence time 40,000 and 1,200 generations ago, respectively.

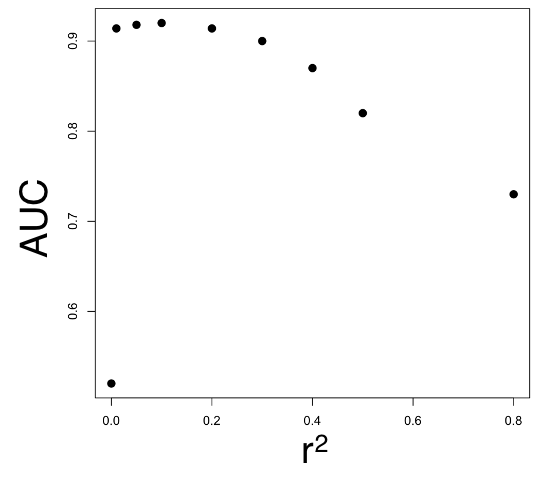

**Figure S5.** AUC values of the ROC analyses, varying the *r^2^* threshold of the *AMI* from 0 to 0.8 in the detection process of ancestry-derived variants.

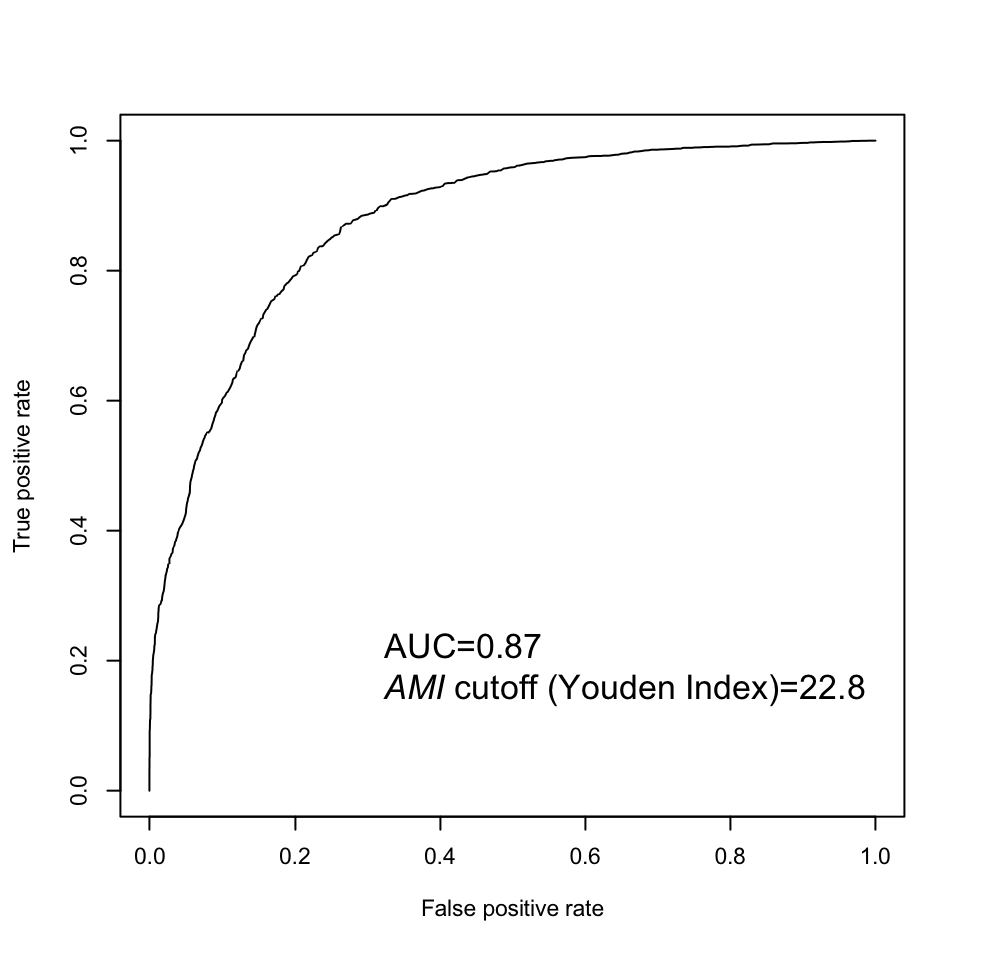

**Figure S6. (A)** ROC curve illustrating the performance of the *AMI* for the detection of the Jomon-derived variants. The ROC curve was drawn based on the simulation of 100 replicates of 1 Mb with the divergence time changed to 800 generations ago (20,000 years ago). The *AMI* showed high accuracy (AUC = 0.87) for discriminating the Jomon-derived variants (type 1) from the other variants (types 2 and 3).

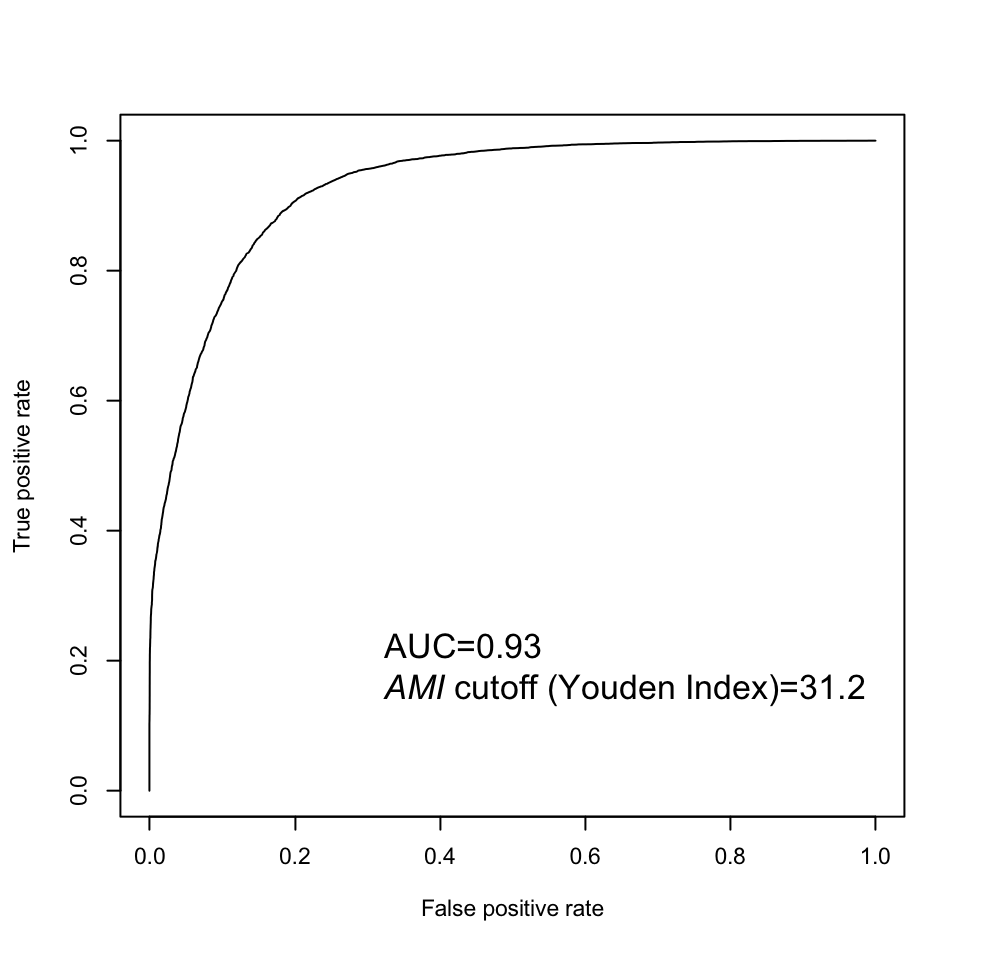

**Figure S6. (B)** ROC curve illustrating the performance of the *AMI* for the detection of the Jomon-derived variants. The ROC curve was drawn based on the simulation of 100 replicates of 1 Mb with the divergence time changed to 1,600 generations ago (40,000 years ago). The *AMI* showed high accuracy (AUC = 0.93) for discriminating the Jomon-derived variants (type 1) from the other variants (types 2 and 3).

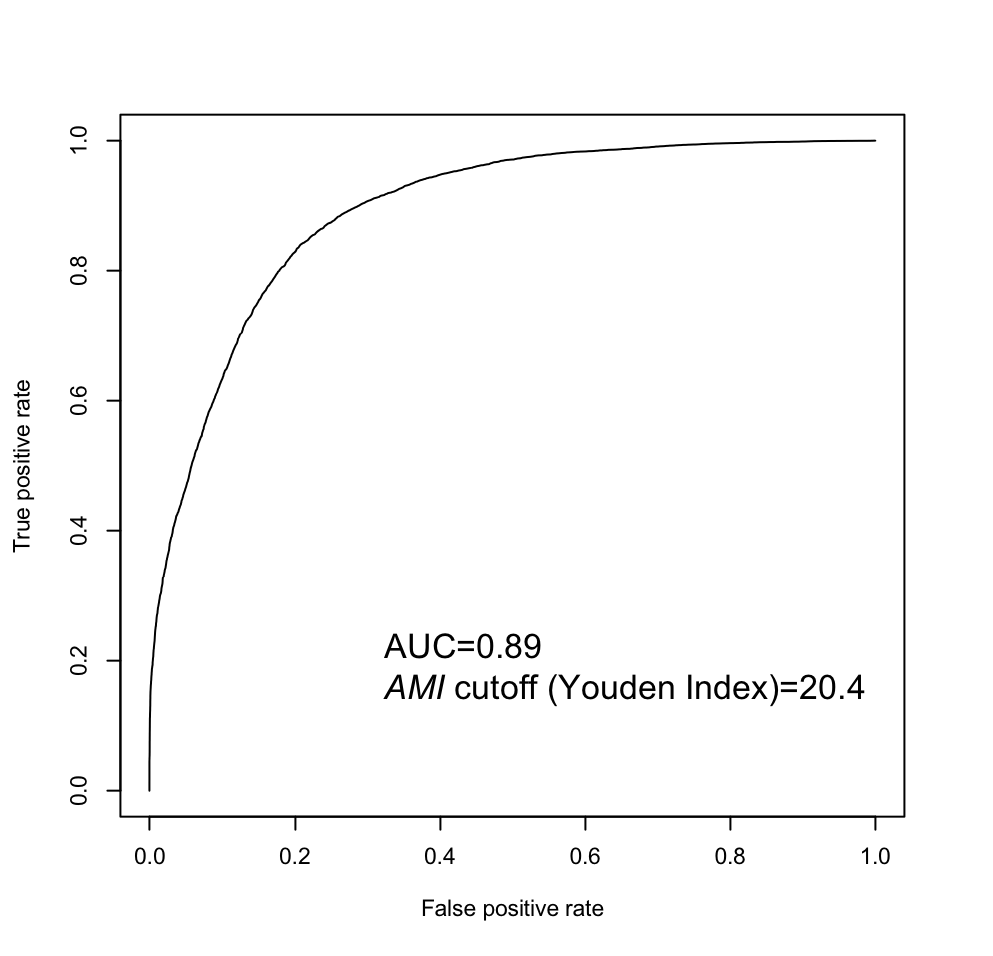

**Figure S6. (C)** ROC curve illustrating the performance of the *AMI* for the detection of the Jomon-derived variants. The ROC curve was drawn based on the simulation of 100 replicates of 1 Mb with the effective population size changed to 10,000. The *AMI* showed high accuracy (AUC = 0.89) for discriminating the Jomon-derived variants (type 1) from the other variants (types 2 and 3).

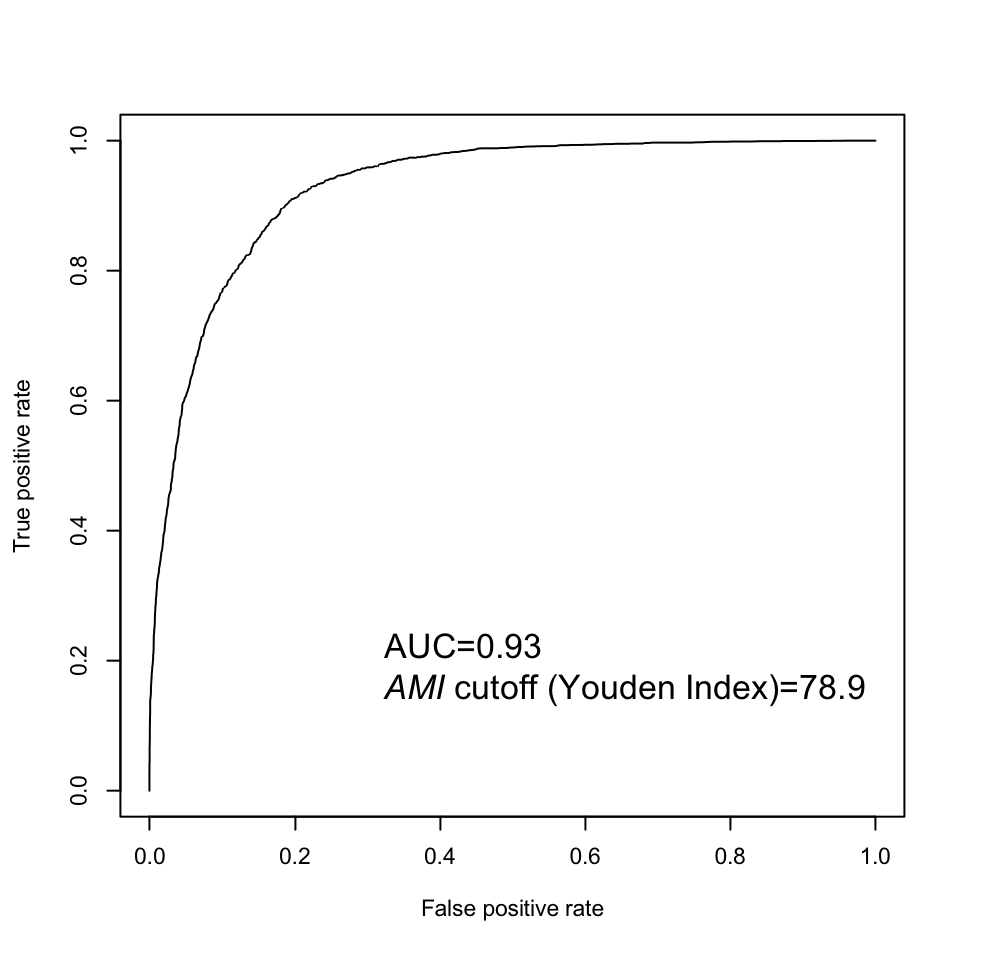
**Figure S6. (D)** ROC curve illustrating the performance of the *AMI* for the detection of the Jomon-derived variants. The ROC curve was drawn based on the simulation of 100 replicates of 1 Mb with the effective population size changed to 1,000. The *AMI* showed high accuracy (AUC = 0.93) for discriminating the Jomon-derived variants (type 1) from the other variants (types 2 and 3).

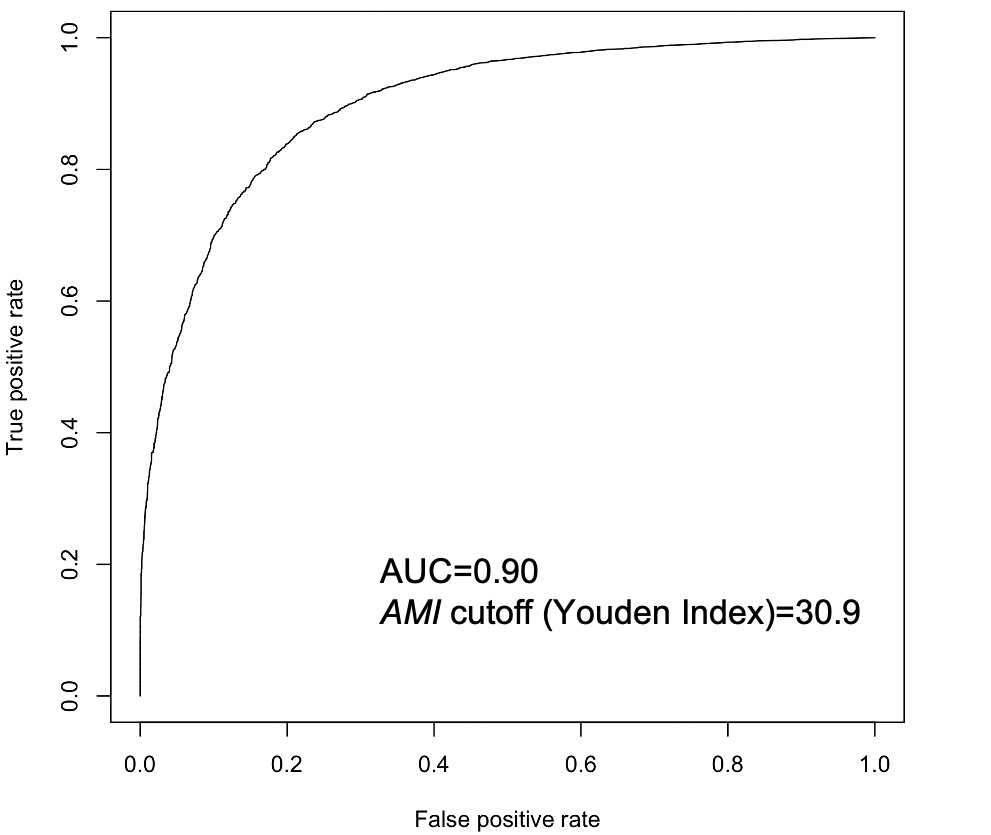

**Figure S6. (E)** ROC curve illustrating the performance of the *AMI* for the detection of the Jomon-derived variants. The ROC curve was drawn based on the simulation of 100 replicates of 1 Mb with the Jomon ancestry proportion of current Japanese population changed to 20%. The *AMI* showed high accuracy (AUC = 0.90) for discriminating the Jomon-derived variants (type 1) from the other variants (types 2 and 3).

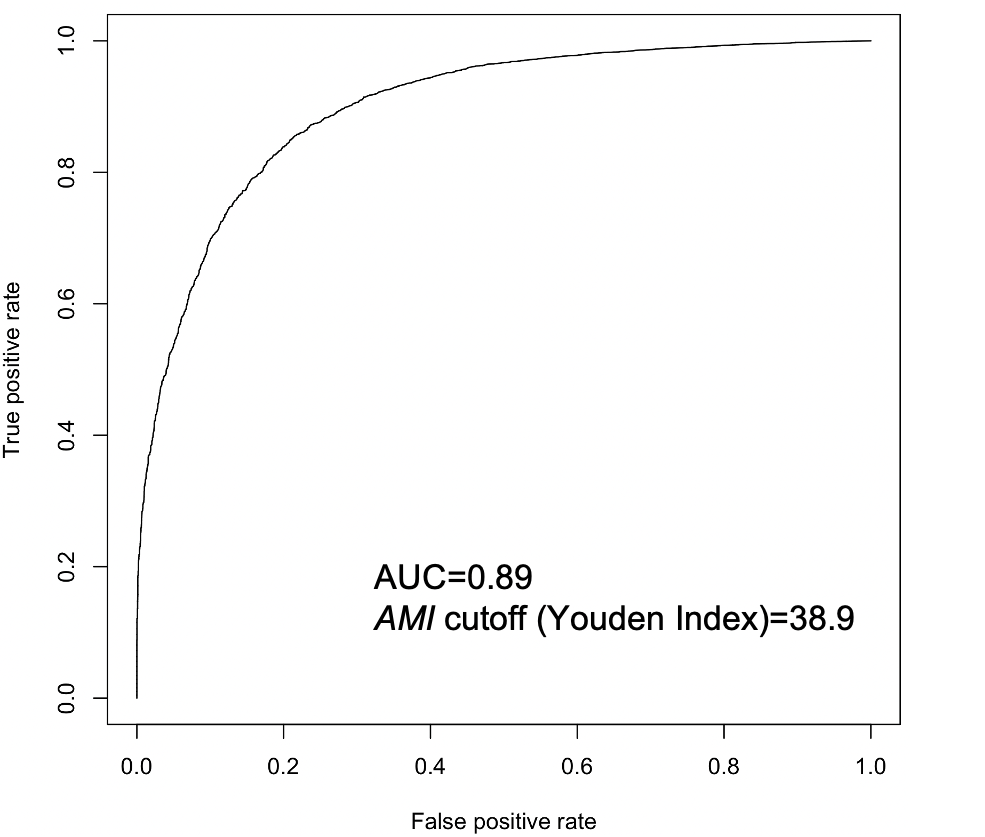

**Figure S6. (F)** ROC curve illustrating the performance of the *AMI* for the detection of the Jomon-derived variants. The ROC curve was drawn based on the simulation of 100 replicates of 1 Mb with the Jomon ancestry proportion of current Japanese population changed to 40%. The *AMI* showed high accuracy (AUC = 0.89) for discriminating the Jomon-derived variants (type 1) from the other variants (types 2 and 3).

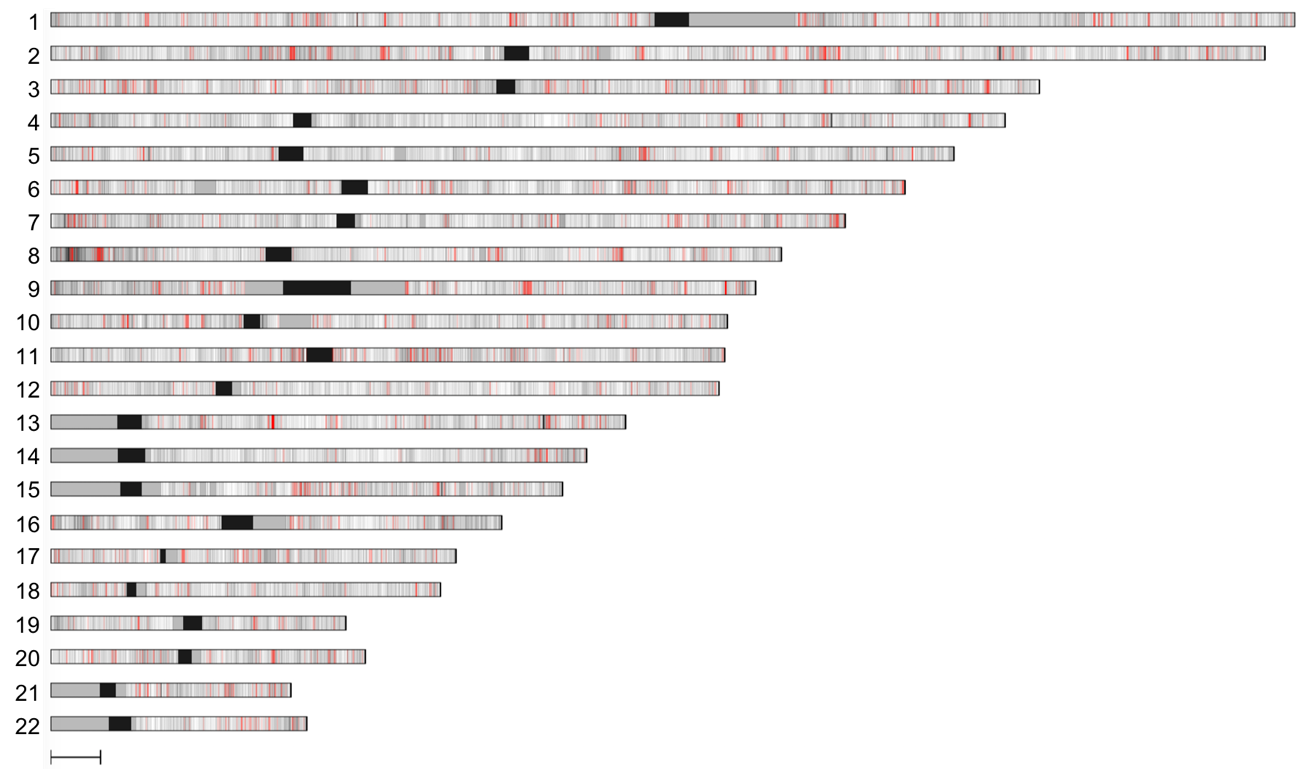

**Figure S7.** Distribution of Jomon-derived variants in the Japanese genomes. The vertical lines represent the Jomon-derived variants, with gray representing SNPs with a Jomon allele frequency of less than 5% and red representing SNPs with a Jomon allele frequency of more than 5%. The black square shows the centromere. The gray square shows the regions with a density of Japanese specific variants below a mean - 1sd of each chromosome (referred to in “Detection of the Jomon derived variants in real data” of the Materials and Methods section). The 10 Mb scale bar is shown under chromosome 22.

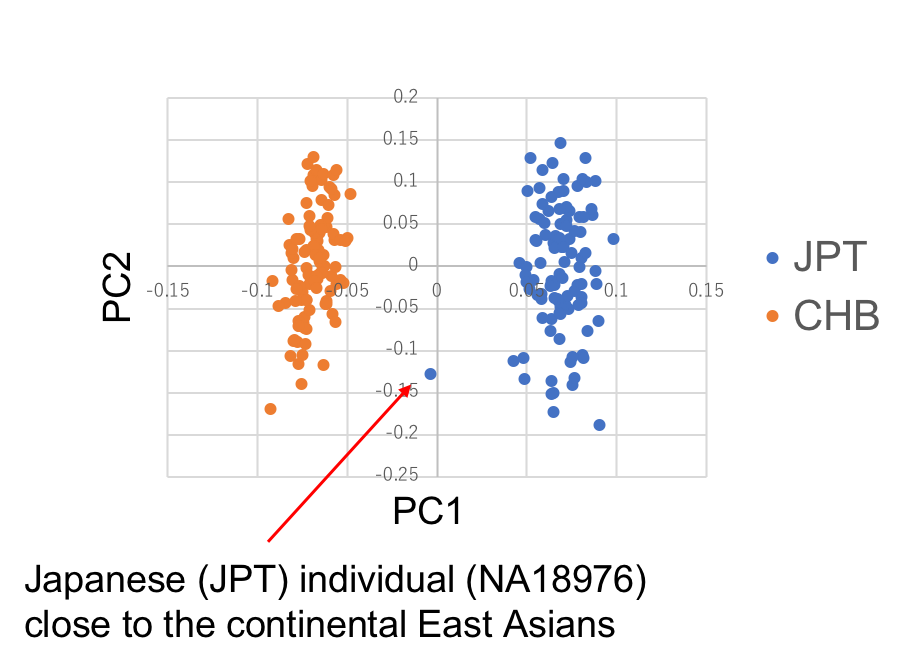

**Figure S8.** Principal component analysis of JPT and CHB of 1000 Genomes project phase III. 104 individuals of JPT (Japanese in Tokyo, Japan) and 103 individuals of CHB (Han Chinese in Beijing, China) are used in the analysis.

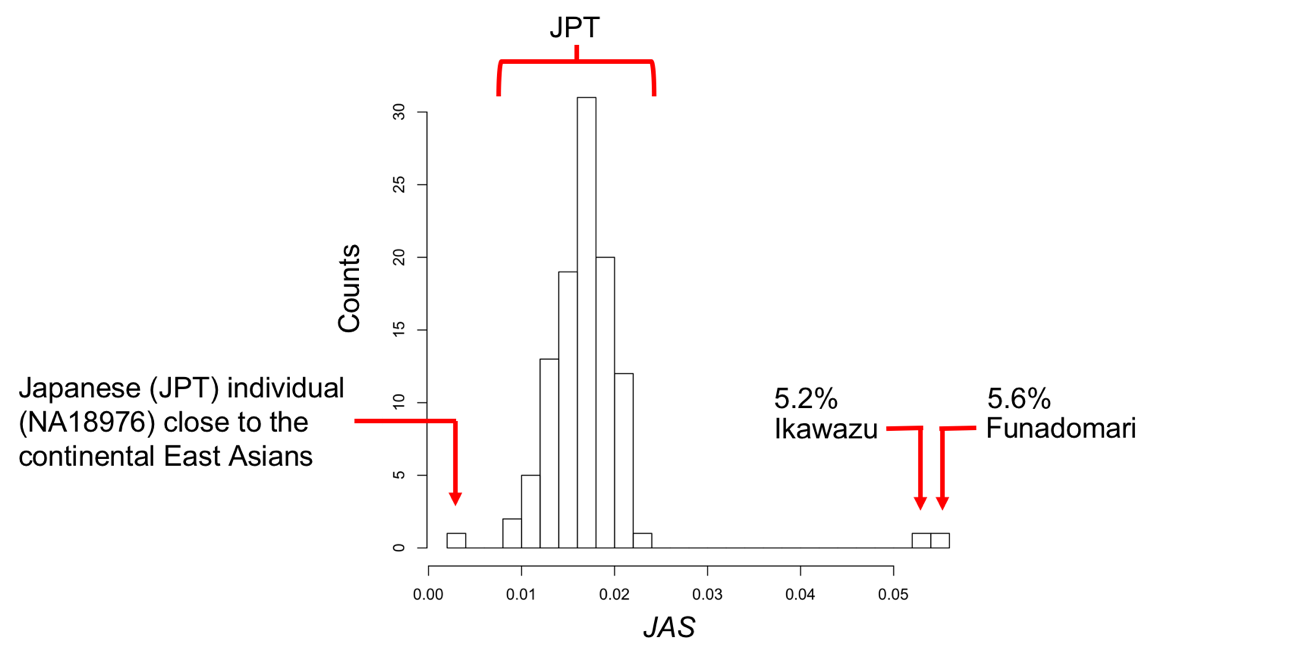

**Figure S9.** Distribution of the *JAS* of the Mainland Japanese and Jomon individuals. The *JAS* were calculated for 104 individuals of JPT (Japanese in Tokyo, Japan) and two Jomon samples (Funadomari and Ikawazu).

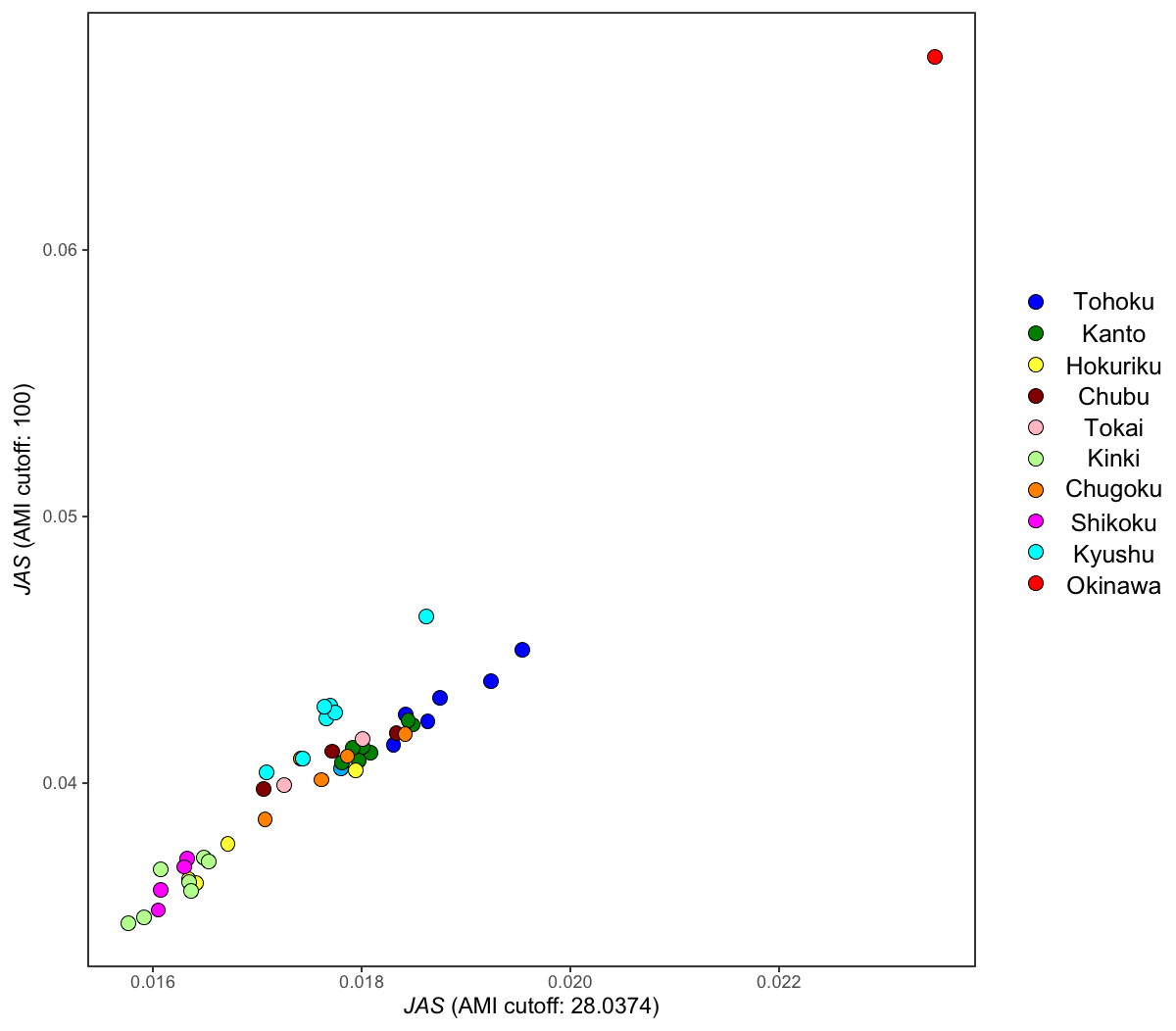

**Figure S10.** Relationship between *JASs* in different cutoff value of the *AMI* in each prefecture. Each prefecture was colored according to the region of Japan in Figure S1. Horizontal axes: *JAS of AMI* cutoff = 28.0374, vertical axes: *JAS of AMI* cutoff = 100

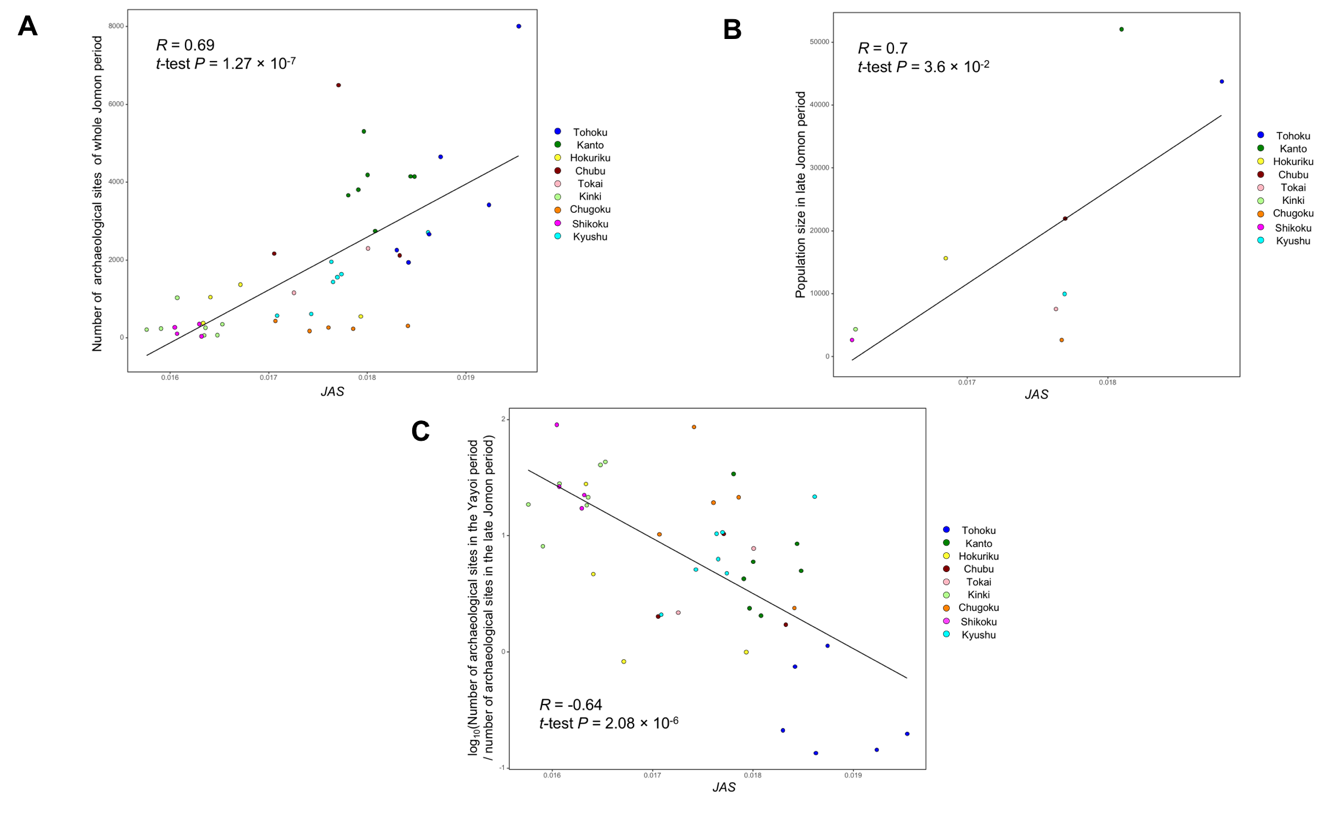

**Figure S11.** Relationship between the *JAS* and values associated with the population size of prefectures in the Jomon to Yayoi periods. The horizontal axis shows the average *JAS* and the vertical axis shows the (A) number of archaeological sites of the whole Jomon period, (B) population size in the Late Jomon period, and (C) log_10_ (number of archaeological sites in the Yayoi period/number of archaeological sites in the Late Jomon period). Pearson's correlation coefficients (*R*), *P* values are shown in each figure. Each prefecture is colored according to the region in Figure S1.

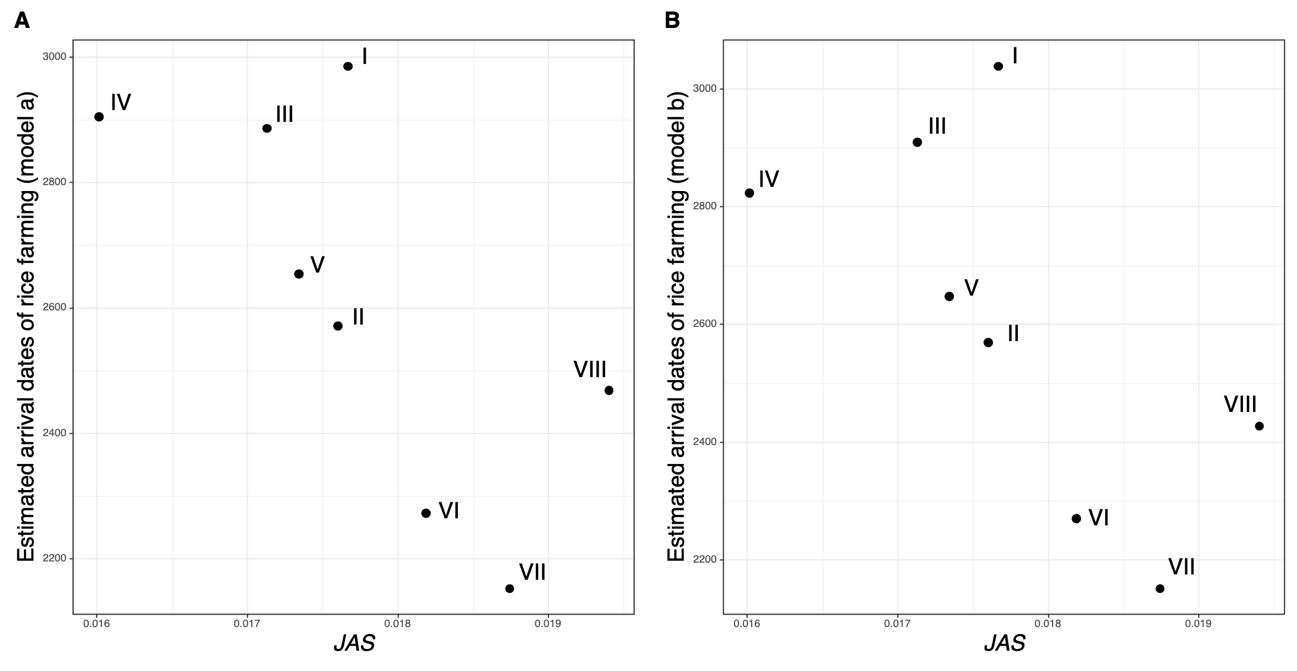

**Figure S12.** Relationship between the *JAS* and arrival dates of rice farming estimated in Crema et al.^8^. Crema et al. estimated the timing of rice farming arrival based on radiocarbon dating of charred rice remains by constructing two different models a and b. We adopt the regional classification of mainland Japan in Crema et al. (Ⅰ to Ⅷ) rather than that of the other analyses in this study. The horizontal axis of (A) and (B) is the *JAS* value in each region while the vertical axis is the arrival dates estimated on model a and b, respectively.

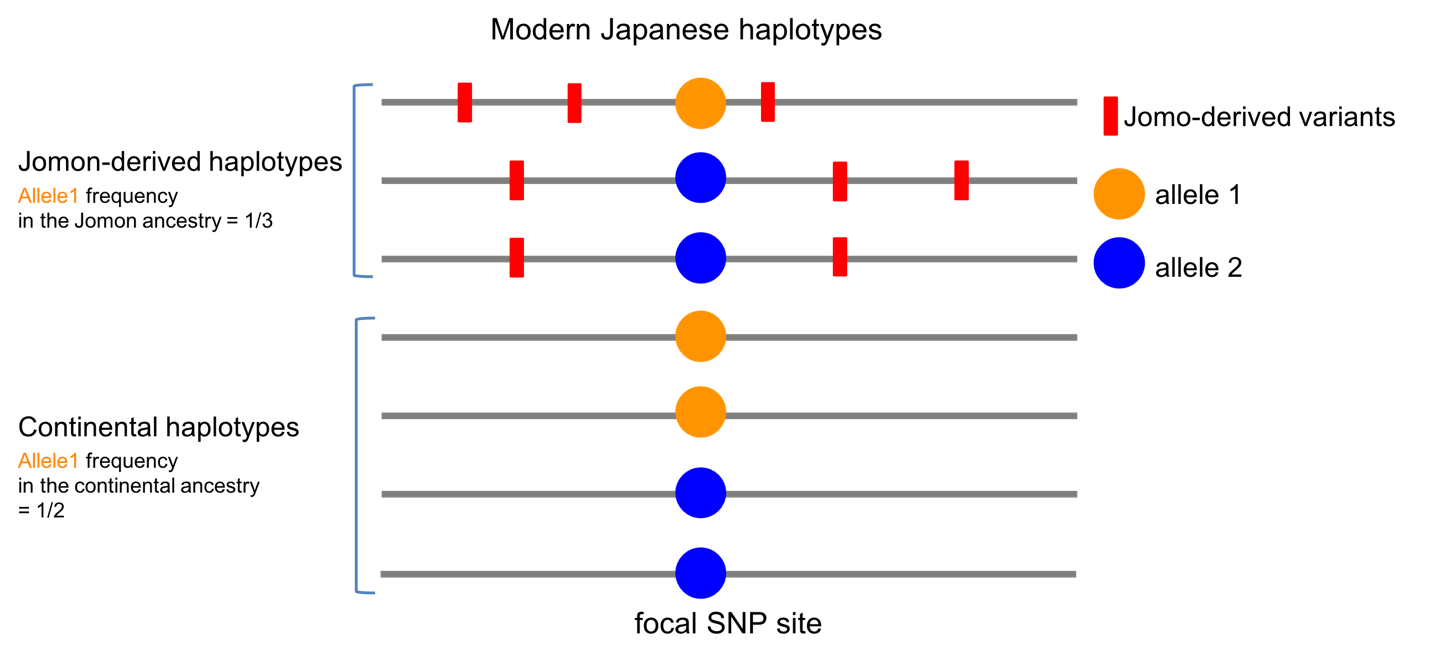

**Figure S13.** Estimation method of the allele frequencies of genome-wide SNPs in the Jomon and continental ancestries of modern Japanese. The modern Japanese haplotypes surrounding a focal variant site can be classified into "Jomon-derived haplotypes" and "continental haplotypes" according to the presence Jomon alleles of Jomon-derived variants (red bar). The allele frequency of the Jomon people in the focal variant site can be estimated by the proportion of each allele within Jomon-derived haplotypes.

**Supplemental Tables**

**Table S1.** The number of samples from each prefecture in Japan.

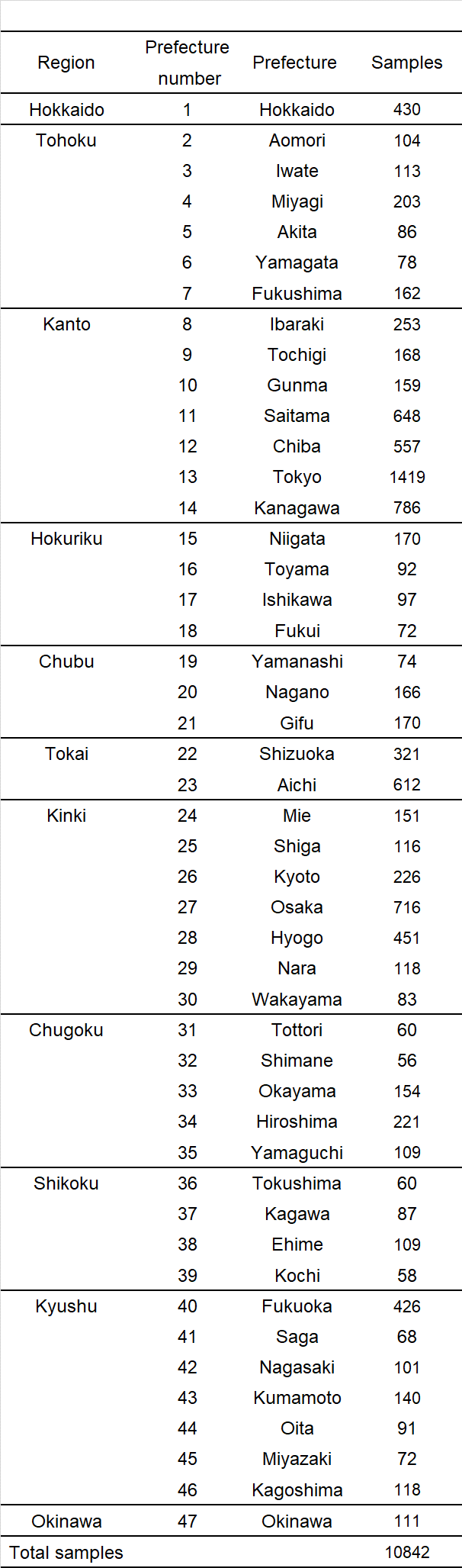

**Table S2.** The Jomon allele score of each region in Japan.

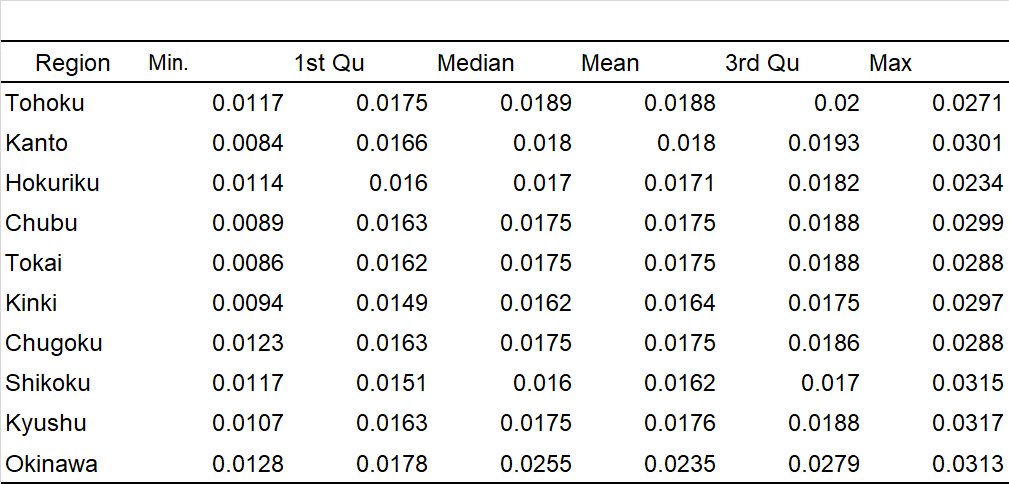

**Table S3.** The Jomon allele score of each prefecture in Japan.

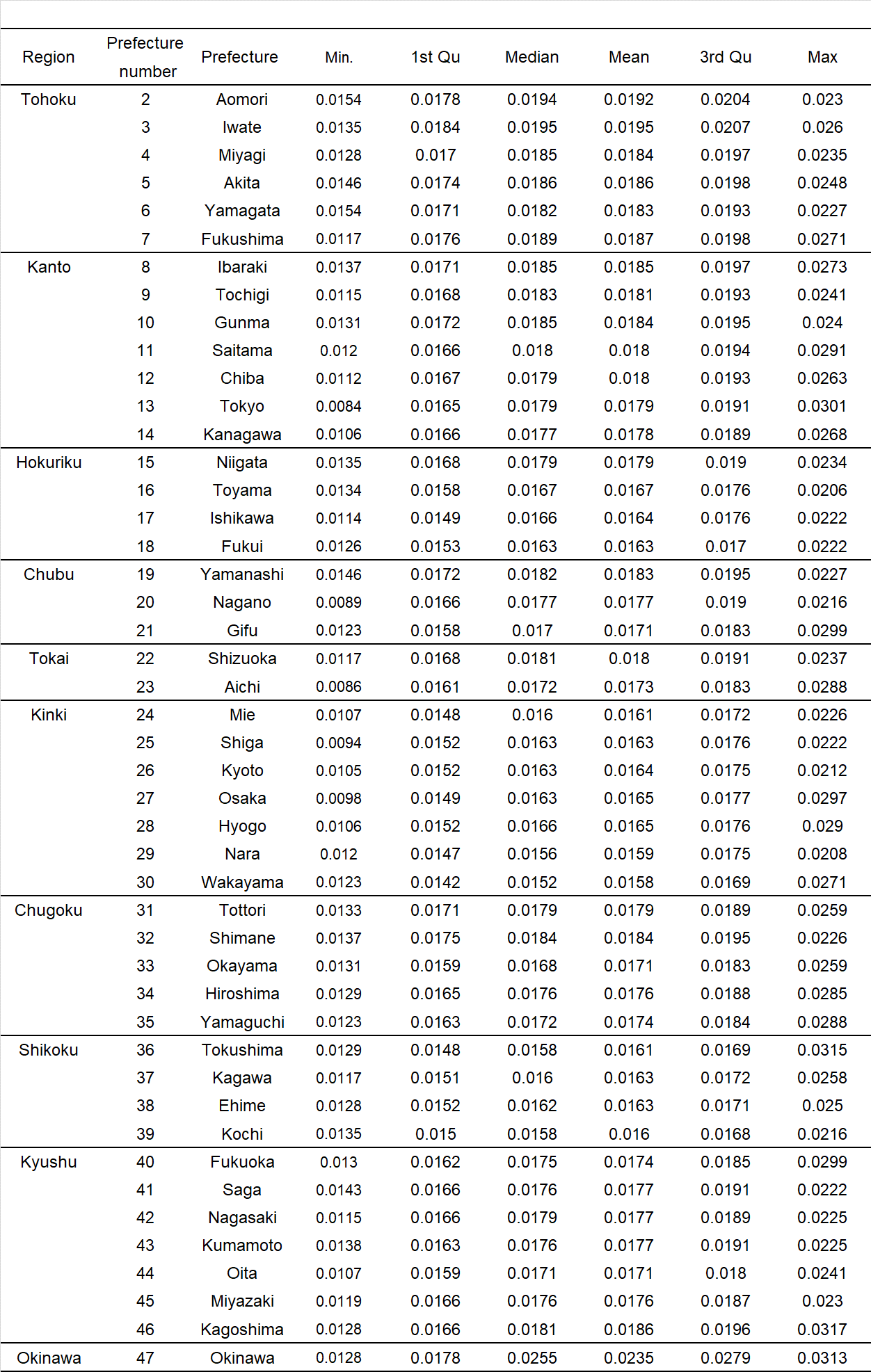

**Table S4** Details in subjected QTL GWAS traits in the current study.

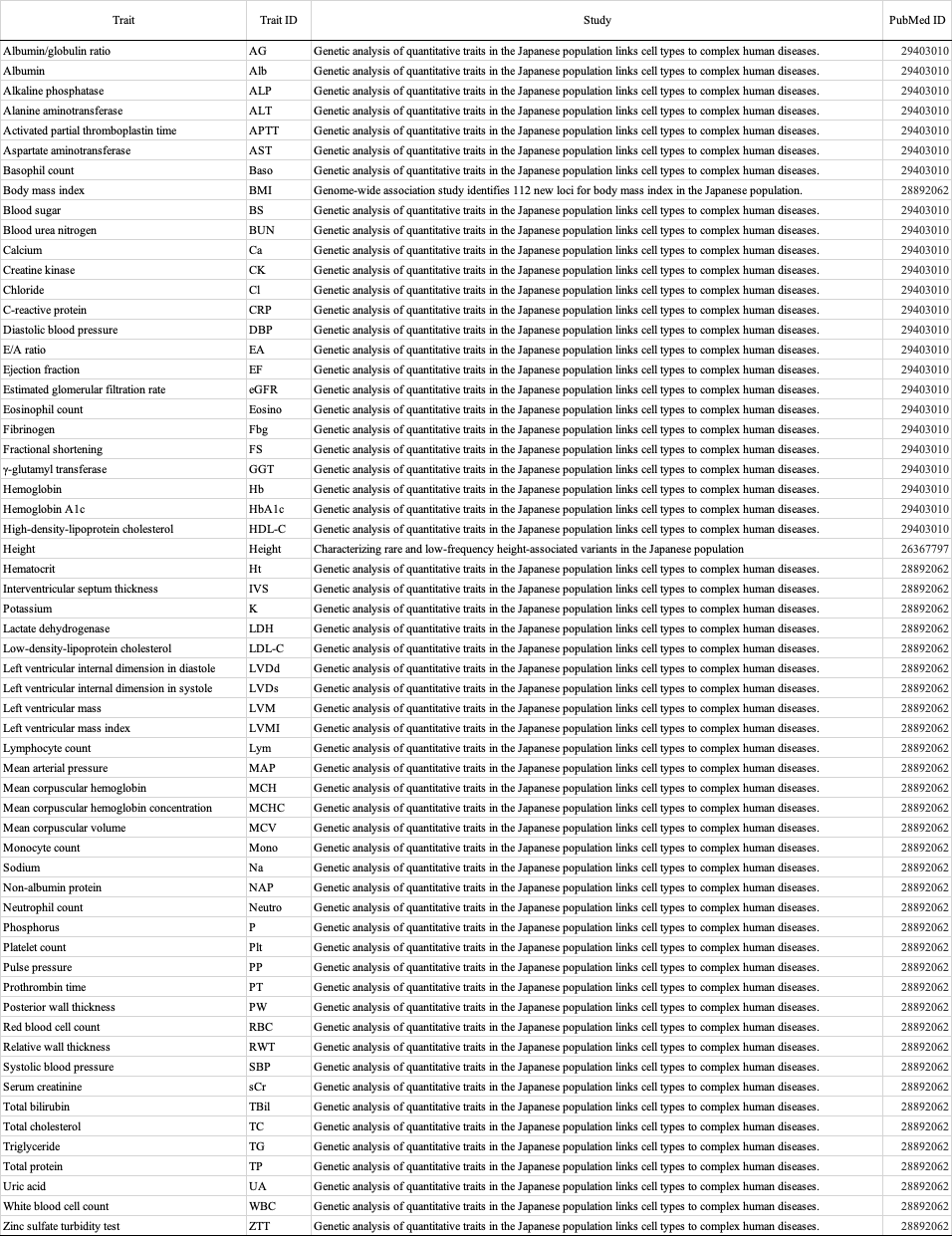

**Table S5** Mean $2\beta f$ value of THC modern Japanese, THC Jomon, and THC continental ancestries and *D* value for each trait, varying the GWAS P value threshold.

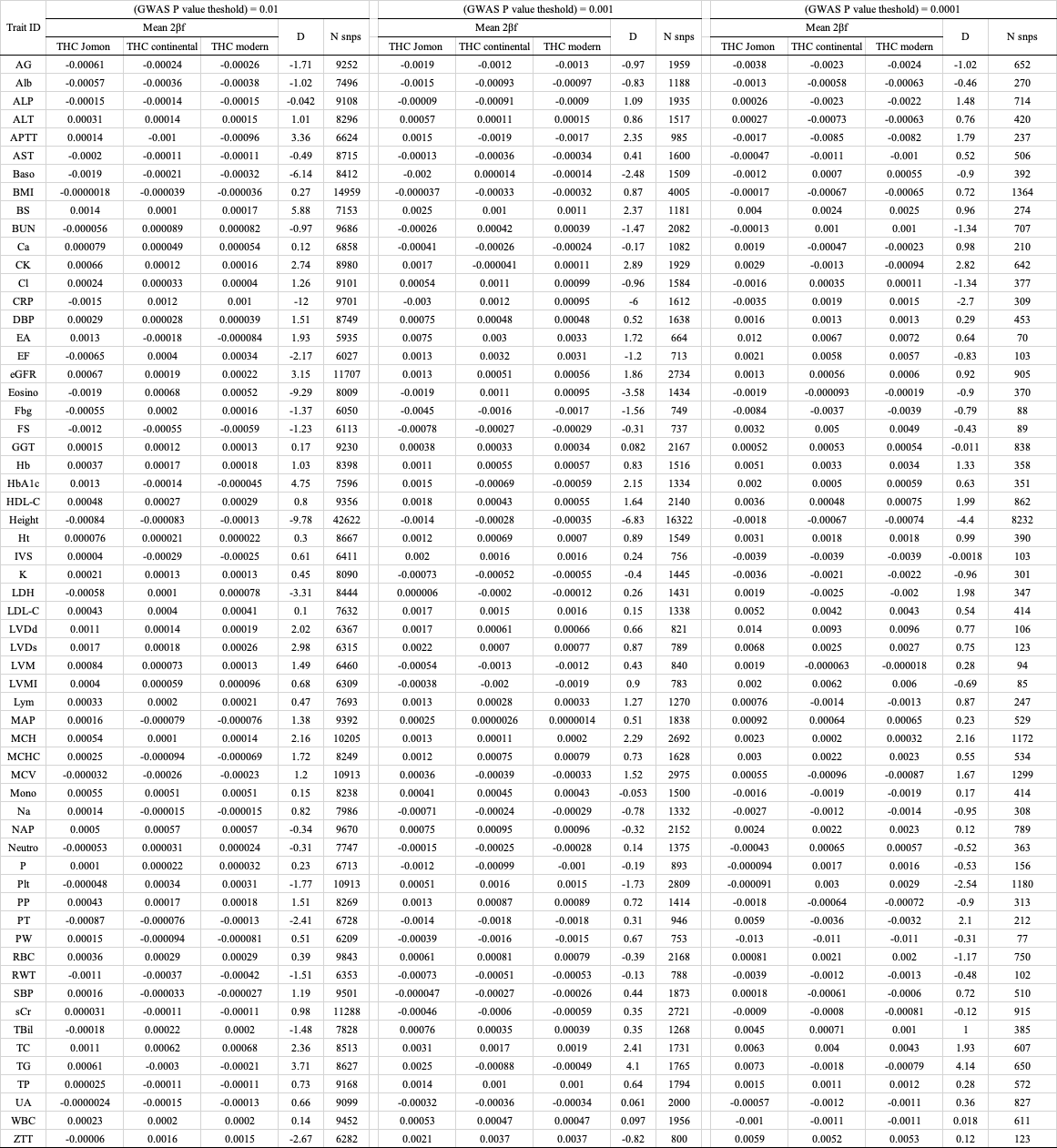
